# HyperTraPS-CT: Inference and prediction for accumulation pathways with flexible data and model structures

**DOI:** 10.1101/2024.03.07.583841

**Authors:** Olav N. L. Aga, Morten Brun, Kazeem A. Dauda, Ramon Diaz-Uriarte, Konstantinos Giannakis, Iain G. Johnston

## Abstract

Accumulation processes, where many potentially coupled features are acquired over time, occur throughout the sciences, from evolutionary biology to disease progression, and particularly in the study of cancer progression. Existing methods for learning the dynamics of such systems typically assume limited (often pairwise) relationships between feature subsets, cross-sectional or untimed observations, small feature sets, or discrete orderings of events. Here we introduce HyperTraPS-CT (Hypercubic Transition Path Sampling in Continuous Time) to compute posterior distributions on continuous-time dynamics of many, arbitrarily coupled, traits in unrestricted state spaces, accounting for uncertainty in observations and their timings. We demonstrate the capacity of HyperTraPS-CT to deal with cross-sectional, longitudinal, and phylogenetic data, which may have no, uncertain, or precisely specified sampling times. HyperTraPS-CT allows positive and negative interactions between arbitrary subsets of features (not limited to pairwise interactions), supporting Bayesian and maximum-likelihood inference approaches to identify these interactions, consequent pathways, and predictions of future and unobserved features. We also introduce a range of visualisations for the inferred outputs of these processes and demonstrate model selection and regularisation for feature interactions. We apply this approach to case studies on the accumulation of mutations in cancer progression and the acquisition of anti-microbial resistance genes in tuberculosis, demonstrating its flexibility and capacity to produce predictions aligned with applied priorities.

## Introduction

Many important processes in the biological, medical, and physical sciences can be classed as ‘accumulation processes’ (Diaz-Uriarte and Herrera-Nieto, 2022; Diaz-Uriarte, 2023), involving the serial stochastic acquisition or loss of discrete traits over time – from evolutionary dynamics (O’Meara, 2012; Johnston and Williams, 2016; Williams et al., 2013) to disease progression (Schill et al., 2024; Beerenwinkel et al., 2015; Colijn et al., 2017). Traits (also called characters, particularly in the evolutionary literature) in this context typically mark the presence or absence of a feature of interest – for example, a particular mutation in a cancer patient (Diaz-Uriarte, 2023; Schill et al., 2024), a given gene lost by a species (Johnston and Williams, 2016), or a given disease symptom presented by a patient (Johnston et al., 2019). Reconstructing the dynamics by which these processes occur can inform our knowledge of the underlying mechanisms (Johnston and Williams, 2016; Williams et al., 2013), make predictions about unmeasured traits and the likely future behaviour of systems in a known state (Diaz-Colunga and Diaz-Uriarte, 2021; Diaz-Uriarte and Vasallo, 2019; Luo et al., 2023; Williams et al., 2013; Colijn et al., 2017), and identify features of the system which determine (or are determined by) progress through accumulation pathways (Johnston et al., 2019; Johnston and Røyrvik, 2020).

The study of the evolution of traits across phylogenetically related lineages has an extensive history and associated literature, but general approaches suitable for large sets of data and features remain challenging. Methods to infer phylogenetic trees, to infer evolutionary dynamics of characters (traits) on phylogenies, and to jointly infer both have been developed (reviewed, for example, in Revell and Harmon (2022) and O’Meara (2012) and included in famous software packages like *phytools* (Revell, 2012) and *corHMM* (Boyko and Beaulieu, 2021)). Another branch of the scientific literature, which remains surprisingly disconnected from the evolutionary picture, focusses on inferring accumulation dynamics in cancer progression (recently reviewed in Diaz-Uriarte (2023) and Schill et al. (2024)). These approaches attempt to describe the acquisition of coupled traits – usually mutations – with time, with classic approaches including Markov modelling of transitions between discrete states with disease observations (Hjelm et al., 2006). Inference methods based on Bayesian networks have played a particularly pronounced role in this field (Beerenwinkel et al., 2015; Ramazzotti et al., 2019; Loohuis et al., 2014; Ramazzotti et al., 2015; Ross and Markowetz, 2016; Montazeri et al., 2016; Szabo and Boucher, 2002). Traditionally (but not exclusively), the cancer literature considered independent, cross-sectional observations of a limited number of binary features, constraining the possible interactions between traits to, for example, pairwise positive influences (but see below).

In an attempt to relax these restrictions, and to support cross-sectional, phylogenetic, and/or longitudinal observations, ‘hypercubic transition path sampling’ or HyperTraPS was developed (Johnston and Williams, 2016; Greenbury et al., 2020). HyperTraPS requires no assumptions about restricted states or independence of feature acquisitions, and is naturally embedded in a Bayesian framework supporting prior information and uncertainty quantification. A focus on transitions between states as the fundamental observation type, rather than individual observations, means that HyperTraPS has always supported data embedded in trees and/or longitudinal sequences, as well as cross-sectional samples. Polynomial rather than exponential scaling in the number of features, and at most linear scaling in the number of observations, permits the efficient analysis of large sets of coupled features where other approaches struggle (Greenbury et al., 2020; Johnston and Williams, 2016). This efficiency arises from a key feature of the approach: an algorithm conceptually similar to transition interface sampling or forward flux sampling in statistical physics (Allen et al., 2009), focussing only on those pathways likely to correspond to observed transitions. While pairwise positive and negative interactions (involving *L*^2^ parameters) are the default target of inference in HyperTraPS, regularisation and model selection approaches support a choice between different parameterisation structures for a given dataset (Greenbury et al., 2020). Prior to and following HyperTraPS, two related approaches – phenotypic landscape inference (Williams et al., 2013) and HyperHMM (hypercubic hidden Markov modelling, Moen and Johnston (2023)) – expanded the parameter space of accumulation models to allow arbitrary interactions between sets of features (not just pairwise interactions), so that, for example, a combination of traits *A, B* can influence trait *C* differently from the combined influence of *A* and *B*. HyperTraPS and these aligned approaches have been used to explore cancer progression, identifying new pathways and interactions (Greenbury et al., 2020; Moen and Johnston, 2023), but their flexibility has also allowed their application in other fields including the evolution of genomes (Johnston and Williams, 2016; Allen and Martin, 2016); multi-drug resistance in tuberculosis (Greenbury et al., 2020; Garcia Pascual et al., 2024); photosynthetic pathways (Williams et al., 2013; Samal and Martin, 2013), and tool-use behaviour (Johnston and Røyrvik, 2020); disease progression in severe malaria (Johnston et al., 2019); and the behaviour of students in online learning (Peach et al., 2021). Predictions from these inferred models about unobserved features and future behaviours have been validated either both using withheld data (Johnston et al., 2019) and independent experiments (Williams et al., 2013). HyperTraPS is certainly not alone in this breadth of application; other approaches from accumulation modelling have also been expanded into other applied fields, notably in the study of HIV drug resistance (Beerenwinkel and Drton, 2007; Posada-Céspedes et al., 2021).

Independently of HyperTraPS, Mutual Hazard Networks (MHN) has been developed more recently (Schill et al., 2020, 2024), driving the cancer accumulation field forward – with other powerful advances including those based on Bayesian networks (Nicol et al., 2021; Diaz-Uriarte and Herrera-Nieto, 2022; Diaz-Uriarte, 2023; Angaroni et al., 2022), permutation analysis (Peterson and Kovyrshina, 2017; Zhang and Wang, 2018), and accounting for tumour ‘phylogenetics’ (reviewed in Schwartz and Schäffer (2017)). MHN uses the same principle as the pairwise, *L*^2^, parameterisation of HyperTraPS to support pairwise positive and negative influences between traits (but does not support higher-order interactions). MHN has recently been embedded in a framework, TreeMHN, allowing observations to be connected in a tree (Luo et al., 2023), matching the native capacity of HyperTraPS to deal with phylogenetically embedded and longitudinal data. To deal with larger sets of features, TreeMHN introduced a sampling approach to the MHN picture, aligning with HyperTraPS’ sampling system.

One difference between these connected approaches is their picture of time. MHN and TreeMHN, for example, contain an implicit sampling time that sets a continuous timescale for the inferred dynamics; this feature has been a target of refinement and subsequent computational acceleration in Gotovos et al. (2021). HyperTraPS and HyperHMM consider only orderings, not timings, of transitions – originally motivated by the absence of timing information in the evolutionary and disease progression settings under study. In many evolutionary applications (and now cancer progression) of accumulation modelling, however, imperfect timing information does exist, in the form of (uncertain) branch lengths in the associated trees – either corresponding to real time or to amounts of evolutionary change. The capacity to deal with uncertain timings is also useful in cancer and disease progression, where observations are typically bounded by inequalities: changes occur at some unknown time between two delineating observation times.

Here, we develop HyperTraPS-CT, an expansion and generalisation of HyperTraPS that connects with absolute timings, either specified precisely or via a range of possible values, allowing uncertainty in observation times to be naturally included. HyperTraPS-CT retains all the existing strengths of HyperTraPS: scalability (having been used for over 120 pairwise-coupled features in Peach et al. (2021)); flexibility in data source (cross-sectional, longitudinal, phylogenetic); the capacity for regularisation and model selection (Greenbury et al., 2020); and a Bayesian implementation. New components of HyperTraPS-CT include the ability to use precise and/or uncertain timing information; the ability to capture arbitrary positive and negative interactions between sets of (not just pairwise) features (mirroring HyperHMM in Moen and Johnston (2023)); and an expanded set of options for predicting unobserved features in observations and future behaviours. We also introduce a suite of visualisation approaches reporting the output of these inference processes, which can readily be applied to other continuous-time models like MHN (Schill et al., 2020), CBN (Montazeri et al., 2016), and H-ESBCN (Angaroni et al., 2022), with a flexible implementation (via command line and R) allowing efficient use of these advances across platforms.

## Methods

### HyperTraPS-CT: Inferring timed evolutionary pathways on a hypercubic transition network

We will work in a picture where observed states of a system are described by the presence or absence of *L* binary traits. There are in total 2^*L*^ different states (that is, unique patterns of presence or absence) which we will refer to with binary strings of length *L*, with 0 or 1 in the *i*th position respectively corresponding to the absence or presence of the *i*th trait (Fig. 1A). We allow Poissonian transitions between states of the system with a characteristic rate: the rate of a transition from *s*_1_ to *s*_2_ is 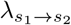 . In monotonic accumulation, where dynamics proceed by individual changes of one trait at a time, 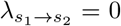 for all *s*_2_ that do not differ from *s*_1_ by an acquisition of exactly one trait. For example, for *L* = 3, *λ*_000*→*001_ may be nonzero, but *λ*_000*→*011_ = 0.

**Figure 1:**
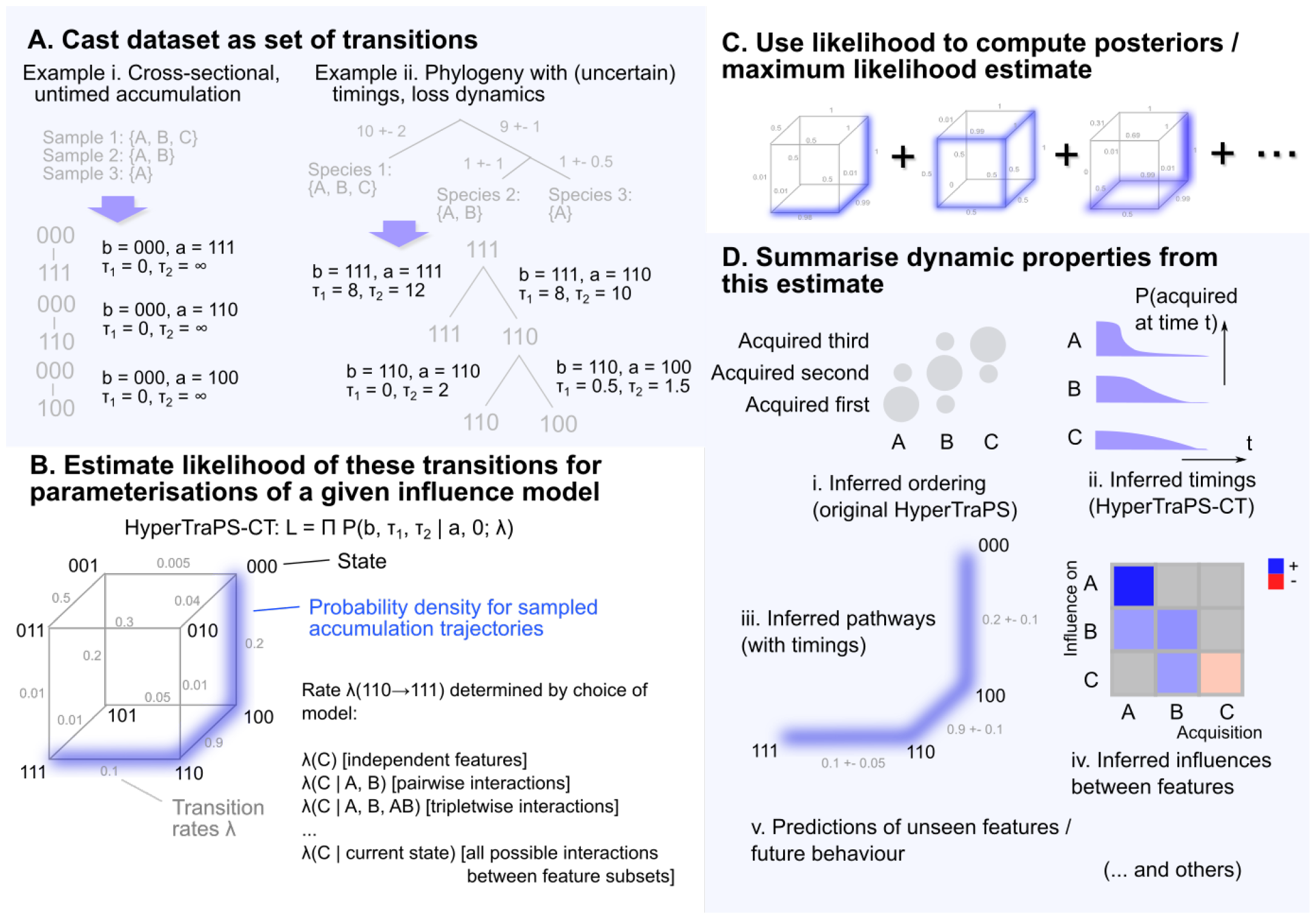
Outline of HyperTraPS-CT approach. **(A)** The relationship between observations is used to create a set of observed transitions. If observations are independent (i) – for example, individual patients, or independent lineages – they can be treated individually by imposing an initial state. If they are longitudinally or phylogenetically related (ii), transitions are inferred from the phylogeny with associated precise timings or uncertain time windows, or with completely unspecified timings where no timing information is available. **(B)** A hypercubic transition network is used to describe evolutionary pathways; Eqn. 1 can be used to estimate, using sampling, a likelihood of observations given a parameterisation of this network *λ*. **(C)** Different parameterisations are explored; for a Bayesian analysis, MCMC and any prior information is used to sample from posterior parameterisations that have a high associated likelihood given the data; for a maximum likelihood estimate the likelihood is optimised. **(D)** The identified parameterisation(s) can then be used to report orderings (i), timescales (ii), pathway structures (iii), influences between features (iv) that are likely given the set of observations, and to make predictions of unseen features and/or future behaviour (v).

Several early instances of HyperTraPS were designed to study systems where *loss* of traits, rather than accumulation, was the driving dynamic (for example, the loss of genes in the reductive evolution of mitochondrial DNA (Johnston and Williams, 2016)). HyperTraPS can readily describe loss dynamics as well as accumulation dynamics, in which case the description above is inverted: 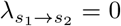 for all *s*_2_ that do not differ from *s*_1_ by a *loss* of exactly one trait. In either case, the network of transitions between states takes a hypercubic structure (Fig. 1B). Evolutionary trajectories of the system are modelled as random walks from some initial state, undergoing transitions randomly according to the rates of available transitions from the current state.

We consider datasets of the form 𝒟 = {*a*_*i*_, *b*_*i*_, *τ*_1*i*_, *τ*_2*i*_}, consisting of a collection of *N* records of ancestral state *a*_*i*_, descendant state *b*_*i*_, and a observation time window (*τ*_1*i*_, *τ*_2*i*_) (Fig. 1A). This window can be used to describe specified, uncertain, or relaxed constraints on observation times (see below). To compute the likelihood of an observation in our dataset, and thus make progress inferring the transition rates that are compatible with observations, we require the probability *P* (*b, τ*_1_, *τ*_2_|*a*, 0; *λ*) that, if a system is in (ancestral) state *a* at time *t* = 0, it will be in (descendant) state *b* at some time between *t* = *τ*_1_ and *t* = *τ*_2_, given a particular set of transition rates *λ*. If it is computationally feasible to analyse all the paths leading from *a* to *b*, this probability can be computed exactly as in MHN (Schill et al., 2020) (see Appendix). However, if we are working with many traits, the number of paths leading from *a* to *b* may be computationally unreasonable to fully sample – particularly if this calculation is inside a loop, for example in a Bayesian search over parameter space. In such cases, we need to employ a sampling scheme that captures the pathways that are most likely. We first define a state *s* as *b*-compatible if *s* has acquired no features that *b* does not have (hence, *b*_*i*_ = 1 for every *i* where *s*_*i*_ = 1; this definition is inverted for loss dynamics). For example, in the case of feature acquisition, 011 is 001-compatible but not 100-compatible. The HyperTraPS algorithm (Johnston and Williams, 2016) gives us a way of constructing paths starting at *a* that are guaranteed to be *b*-compatible, and thus to end at *b*, allowing us to avoid wasting computational time analysing paths that will not correspond to observations.

In Algorithm 1 we present an approach to estimate *P* (*b, τ*_1_, *τ*_2_|*a*, 0; *λ*), which we call HyperTraPS-CT (hypercubic transition path sampling in continuous time). Eqn. 1 gives the central, general expression for the probability estimate (denoted 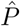). Some special cases merit further attention; we will treat the *λ* dependence as implicit for these. First, if *a* and *b* are identical, the expression reduces to the exact form *P* (*b, τ*_1_, *τ*_2_|*b*, 0) = exp(−*β*_*b*_*τ*_1_) (Step 1 in Algorithm 1), simply describing the probability that the system is still in state *b* after a time window *τ*_1_. Second, if *τ*_1_ = 0 and *τ*_2_ = ∞, we obtain the probability that, given that the system is in state *a* at *t* = 0, *b* is encountered at *any* time in the future. Eqn. 1 then reduces to 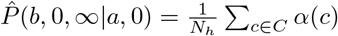, sampling the properties *α*(*c*) of path *c* in the set of paths *C* sampled by *N*_*h*_ independent random walkers (in HyperTraPS(-CT), these random walkers are constrained to follow only *b*-compatible paths – that is, paths that will contribute to the likelihood calculation – and the amount of constraint this entails is recorded and used in the likelihood estimation). This is exactly the quantity reported by the original HyperTraPS algorithm, without considering continuous time. Third, if *τ*_1_ = *τ*_2_ = *τ*, we enforce that the system must be in state *b* at an exact time *τ* after it is observed in state *a*, corresponding to an exactly-specified time window between the two observations.

The derivation of Eqn. 1, Algorithm 1, and these specific results is provided in the Appendix, along with further mathematical details. Briefly, we exploit two facts from the assumption of Poissonian dynamics. First, the ‘arrival time’ – the sum of the transition times through the pathway *c* – follows a hypoexponential distribution, which can be integrated over all times in the window from *τ*_1_ to *τ*_2_. Second, the ‘dwell time’ – the time the system remains at *b* – follows a simple exponential form; accounting for all patterns of arrival and dwell times then characterises the probability of a given observation.

#### Algorithm 1. Hypercubic transition path sampling in continuous time (HyperTraPS-CT). Requires a start state *a*, end state *b*, time window *τ*_1_, *τ*_2_, and transition rates *λ*.

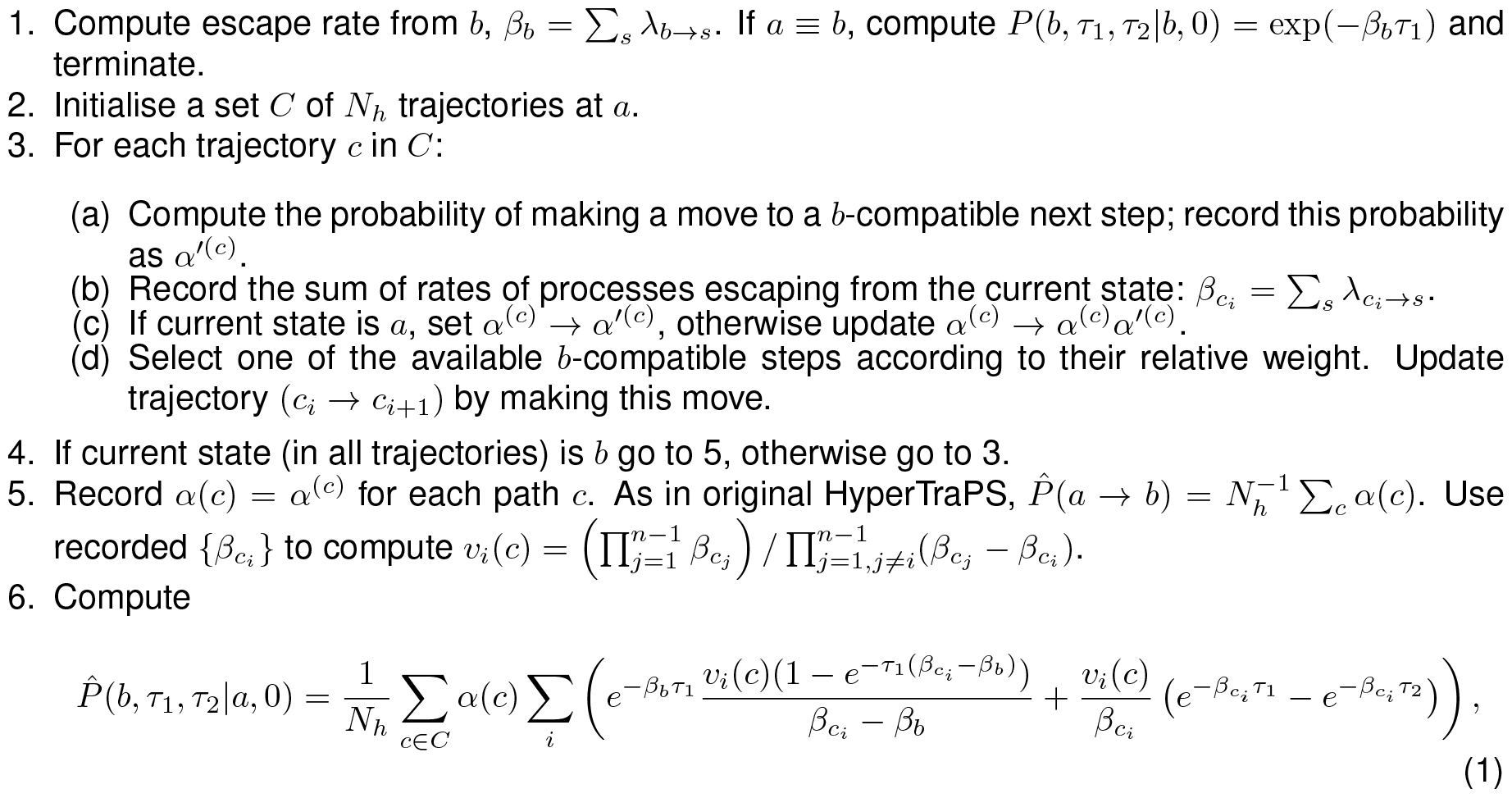

Given the estimated probability from Algorithm 1, we construct an approximate likelihood associated with the dataset 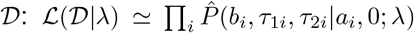. This likelihood function now enables us to perform inference of the transition networks *λ* most compatible with observed data (Fig. 1C). In the Appendix we present a Bayesian MCMC algorithm that produces posterior distributions on transition rates given observations (Fig. 1C-D) and naturally allows prior information on the system to be included in the analysis. We also use a variety of test cases to demonstrate that this approach can infer the true parameterisations of synthetic evolutionary state spaces, and can accurately reconstruct the orderings and timescales of evolutionary events. In addition, we demonstrate how the inclusion of prior information about the evolving system (for example, forbidding some transitions) can be used to increase efficiency and refine the resultant posteriors. All code for these test cases is available at https://github.com/stochasticbiology/hypertraps-ct, with illustrative examples at https://github.com/StochasticBiology/hypertraps-ct/blob/main/docs/hypertraps-demos.pdf.

In Supp. Figs. S1-S2 we demonstrate a set of calculations and verification case studies for the principle of HyperTraPS-CT. The distribution of inferred timescales for an example set of transitions matches the analytic result for Poisson dynamics under different parameterisations (Fig. S2A). The rates on transitions for a model involving a single, simple pathway (Fig. S2B) and a random set of transition rates across multiple pathways (Fig. S2C) are well captured in the inference process, with strong discrepancies only arising for a limited set of rare transitions (Fig. S2D).

## Results

### Basic outputs from HyperTraPS-CT

Fig. 2 gives a first example of output from HyperTraPS-CT, initially using the *L*^2^ parameterisation as also used in mutual hazard networks (MHN, Schill et al. (2020)). Here, the source data is generated from a process supporting two competing accumulation pathways (as used in Greenbury et al. (2020) and Moen and Johnston (2023)). One pathway involves progressive accumulation of feature 1, then 2, then 3, and so on. The other pathway involves progressive accumulation of feature *L*, then *L* − 1, then *L* − 2, and so on. Each progressive accumulation takes 0.1 time units to occur.

**Figure 2:**
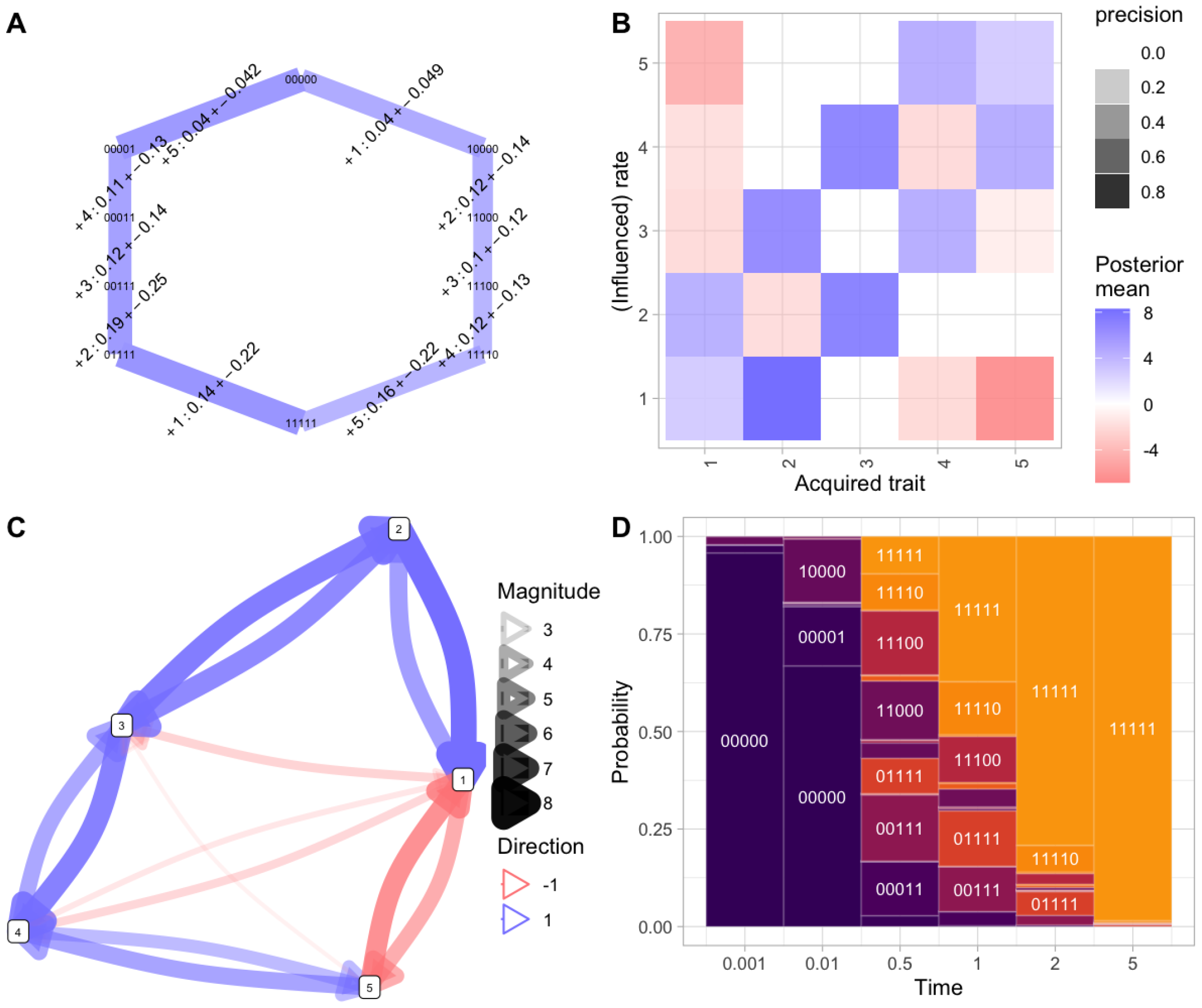
Basic HyperTraPS-CT output for competing pathways. **(A)** Most probable inferred transitions, demonstrating the two competing pathways (left, features 1, 2, 3, …; right, features 5, 4, 3, …). Widths of edges give the probability flux through each edge; edge labels give the feature acquired and the range of associated timings for that transition; node labels give states. **(B)** Following the plot styling in Luo et al. (2023), summary of posterior distributions (mean and coefficient of variation) on the rates of the *L*^2^ parameterisation for this system. Diagonal elements give the base acquisition rates for each feature; off-diagonal elements give the influence of an acquired feature (column) on the acquisition of another feature (row). The opacity of each point reflects its posterior width: ‘precision’ here is max(0, 1 − *CV*) where *CV* is the posterior coefficient of variation. The cross-repression of the first two steps, and promotion of the consequent pathway steps, is clear. **(C)** Inferred network of influences between features, reporting the elements of (B) filtered for low CVs (hence with high posterior probability of being nonzero). **(D)** Probabilities of different states at different snapshot times during system evolution.

Fig. 2 demonstrates some outputs from inference for the *L* = 5 case. Fig. 2A shows transitions through the hypercubic state space inferred to occur with high probability, and their inferred associated timescales. Fig. 2B-C show maps combining the base rates of acquisition of each feature, and how the acquisition of each feature influences the rate of each other feature. Fig. 2B follows the protocol of Luo et al. (2023), representing the *L*^2^ model parameters as a matrix. The Bayesian implementation of HyperTraPS-CT allows uncertainty to be quantified: here, a parameter’s ‘precision’ is reported as max(0, 1 − *CV*) using the coefficient of variation of that parameter’s posterior. Fig. 2C provides a network representation mirroring Fig. 2B, which we can later generalise to the case where collections of features influence each other. Fig. 2D gives predicted states of the system at a collection of ‘snapshot’ observation times. Throughout these plots, the two-pathway structure is clear, including in the specific pathways in Fig. 2, the patterns of base rates and cross-repression between features in Figs. 2B-C, and the predicted states of the system in Fig. 2D.

### Flexible inference with different data and model structures, optimisation, regularisation, and uncertainty

#### Cross-sectional, longitudinal, and phylogenetic datasets

HyperTraPS-CT can naturally handle cross-sectional data (imposing a given ‘ancestral state’ *a*_*i*_ for each *b*_*i*_ observation, for example *a*_*i*_ = 0^*L*^ in the case of accumulation dynamics starting from an initial state with no traits acquired), longitudinal data (decomposed into (*a*_*i*_, *b*_*i*_) pairs through the time course, independent by the Markov property), and phylogenetically embedded data (*a*_*i*_ corresponding to ancestral nodes and *b*_*i*_ to descendant nodes, with each (*a*_*i*_, *b*_*i*_) pair again Markov independent). Fig. 1 illustrates these cases. In this final case, a method for reconstructing ancestral states from modern observations is typically required. If feature acquisitions are assumed to be rare and irreversible events, this process is normally straightforward for accumulation processes: we assume that an ancestor had acquired a feature if all its descendants have it, otherwise we assume that the ancestor did not have the feature and any descendants possessing it acquired it independently (the ancestral state is given by the bitwise AND operator applied over descendants). This picture is readily inverted in the case of loss dynamics, where the bitwise OR operator would apply. For more complex dynamics, tools from phylogenetic reconstruction like the Camin-Sokal (Camin and Sokal, 1965) or Wagner (Eck and Dayhoff, 1966; Kluge and Farris, 1969) methods can be used. In all cases, the (*a, b*) form for an observation – with an associated time window, which may be infinite, wide, or precise (zero-width) – unifies the input data structure.

#### Parameter structures: zero, pairwise, setwise, or arbitrary interactions between features

HyperTraPSCT accepts a range of different parameter structures. The first trivial case, only of applied use as a null hypothesis, is the zero-parameter case, where every feature is acquired independently with the same rate. The next case (independent features) involves a parameter for the ‘base rate’ of acquisition of each feature *F*_*i*_, for a total of *L* parameters. The next case (pairwise interactions) involves these base rates and *L*^2^ − *L* further interaction parameters, describing how the acquisition of feature *F*_*i*_ influences the base rate of feature *F*_*j*_. With a total of *L*^2^ parameters, this setup was introduced by Hjelm et al. (2006), and generalised to allow negative interactions in HyperTraPS (Johnston and Williams, 2016) and published independently as mutual hazard networks (Schill et al., 2020). Unlike those methods, we continue here, with an ‘*L*^3^’ parameterisation allowing pairs of features *F*_*i*_, *F*_*j*_ to have non-additive effects on the acquisition of feature *F*_*k*_; an ‘*L*^4^’ parameterisation allowing triples *F*_*i*_, *F*_*j*_, *F*_*k*_ to have effects independent from their constituent pairs; higher-order models can in principle also be applied. The limiting case is where each of the *L*2^*L−*1^ edges on the hypercube have independent rates, used in HyperHMM (Moen and Johnston, 2023) and corresponding to the case where subsets of features of all possible sizes can have independent influences on a transition.

HyperTraPS-CT allows inference using any of these model choices, with different capacities for interactions between features. The case where every edge on the transition network has an independent weight is likely to correspond to overfitting for reasonable cases. Here, individual parameter values will not all be uniquely identifiable or informative, many different parameterisations may give the same likelihood, and the task more resembles machine learning (finding a parameterisation that generates useful predictions) than parameter inference (identifying interpretable values for given parameters). Conversely, assigning parameters based on individual features alone is likely to underfit data generated by processes where interactions between features are important. Here, parameters will certainly take interpretable values, but may omit important mechanistic information about interactions. The *L*^2^ case risks omitting information about triplet interactions, the *L*^3^ case about quartets, and so on. Model selection and regularisation (described below) can be used to find the optimal parameter structure for the details of a given dataset.

In Fig. 3A we give an example of how different parameter structures omit or capture different mechanisms. The data in this example are generated from a process where pairs exert different influence on feature acquisitions than their constituent individual members. Specifically, the accumulation pathway of the final three features is determined by whether two, or three, of the first three features have been acquired.

**Figure 3:**
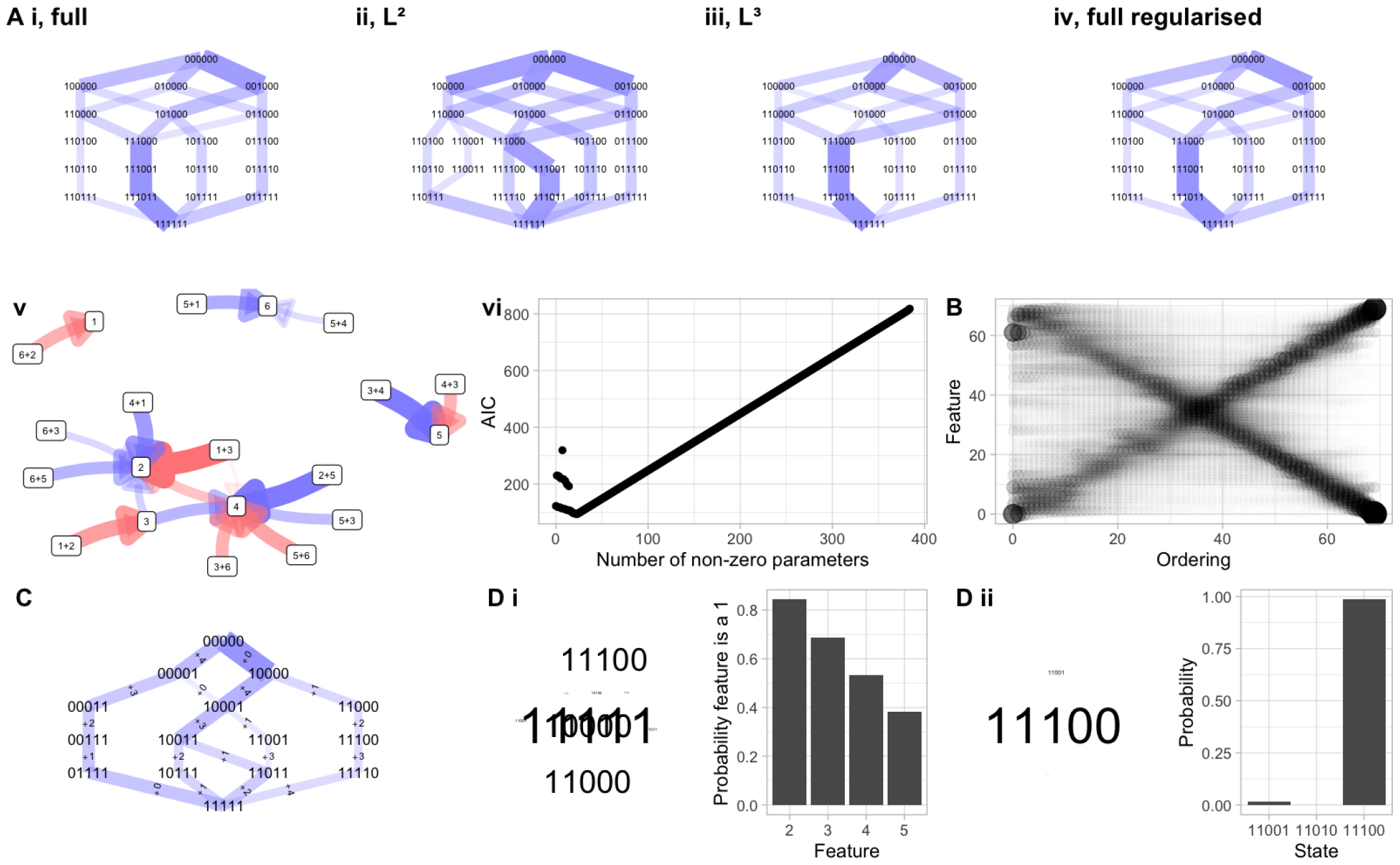
Model structure, scaling, priors, and predictions with HyperTraPS. **(A)** Different parameterisations of the hypercubic transition graph. Here, data are generated from a process where pairs of features influence the acquisition of other features. Inferred transition networks are shown (edge widths give probabilities of transitions) for several models: (i) every edge has an independent parameter; (ii) each feature may have positive or negative influence on the basal rate of each other (as in original HyperTraPS and Mutual Hazard Networks); (iii) each *pair* of features may have additional positive or negative influence on the basal rate of each other feature; (iv) as (i), but following regularisation to remove unimportant edges. In (v) the network of influences between features and pairs of features in the *L*^3^ model (iii) is plotted as in Fig. 2C, showing the effect of higher-order interactions. (vi) shows the progress of the regularisation process in (iv), where the initial 384 parameters are progressively pruned (removing model complexity without sacrificing likelihood) until the information criterion (here AIC) starts to increase as further removal of parameters compromises the likelihood. **(B)** Inference of the competing-pathway model for *L* = 70 features, using 2*L* observations, demonstrating the scalability of HyperTraPS-CT. The same double-pathway structure as before is used; the size of a point reflects the probability that a given feature is acquired in a given step in the accumulation process. **(C)** Inference of the competing-pathway model using priors to enforce an ordering on the basal rates for each feature: 1 *>* 5 ≫ *others*. **(D)** Using learned hypercubes for predictions. (i) Predicting the values of hidden features in the observation 1????. Word cloud gives inferred true states; bar chart gives the probability that each feature takes the value 1. (ii) Predicting the next step from state 11000. Both word cloud and bar chart give inferred next state.

As in Moen and Johnston (2023), the pairwise-interaction picture (*L*^2^ here and in Greenbury et al. (2020); also mutual hazard networks (Schill et al., 2020; Luo et al., 2023; Schill et al., 2024)) cannot capture these higherorder interactions. Comparing Fig. 3Ai with Aii, the *L*^2^ parameterisation is forced to assign nonzero probabilities to a range of pathways that are not present in the generating model, because of the requirements of adjusting lower-order rate parameters to estimate the influence of higher-order processes. The *L*^3^ parameterisation (Fig. 3Aiii) has parameters supporting the nonadditive influence of pairs of features on acquisition rates – as in the generating model – and hence captures the dynamic structure. The corresponding inferred network of interactions, generalising the matrix picture of Fig. 2B, is shown in Fig. 3Av. The all-edges model in Fig. 3Ai is comparable to the target of inference in HyperHMM (Moen and Johnston, 2023); as such a highly-parameterised model will typically reflect overfitting, regularisation can be used to prune extraneous parameters, leaving only those higher-order interactions needed to describe the data (Fig. 3Aiv,vi; see next subsection).

#### Model regularization

In addition to specifying a basic model structure from one of these families, HyperTraPS-CT supports variable selection through regularisation. This can be performed in several ways. First, stepwise parameter removal (as in Greenbury et al. (2020)) where, after a model is fitted, the parameters are progressively set to zero, with the parameter removed at each step being the one that has least influence over the likelihood at that step. The minimum AIC (or other criterion) parameterisation can then be chosen, retaining the set of interactions that are necessary and sufficient to best describe the data (Fig. 3A vi). Second, using a penalised likelihood, where model complexity is included as a penalty in either the maximum-likelihood optimisation or the MCMC process (see below). The penalised complexity can either be the number of nonzero parameters, following an information-criterion-like approach, or the magnitude of parameter values, following a LASSO-like approach. This approach is used in the tuberculosis and cancer case studies below (Figs. 5-4). We generally found penalising the number of nonzero parameters (akin to an AIC-like approach) to give reproducible and robust results, but the best approach (and whether to penalise at all) will in general depend on the scientific question.

**Figure 5:**
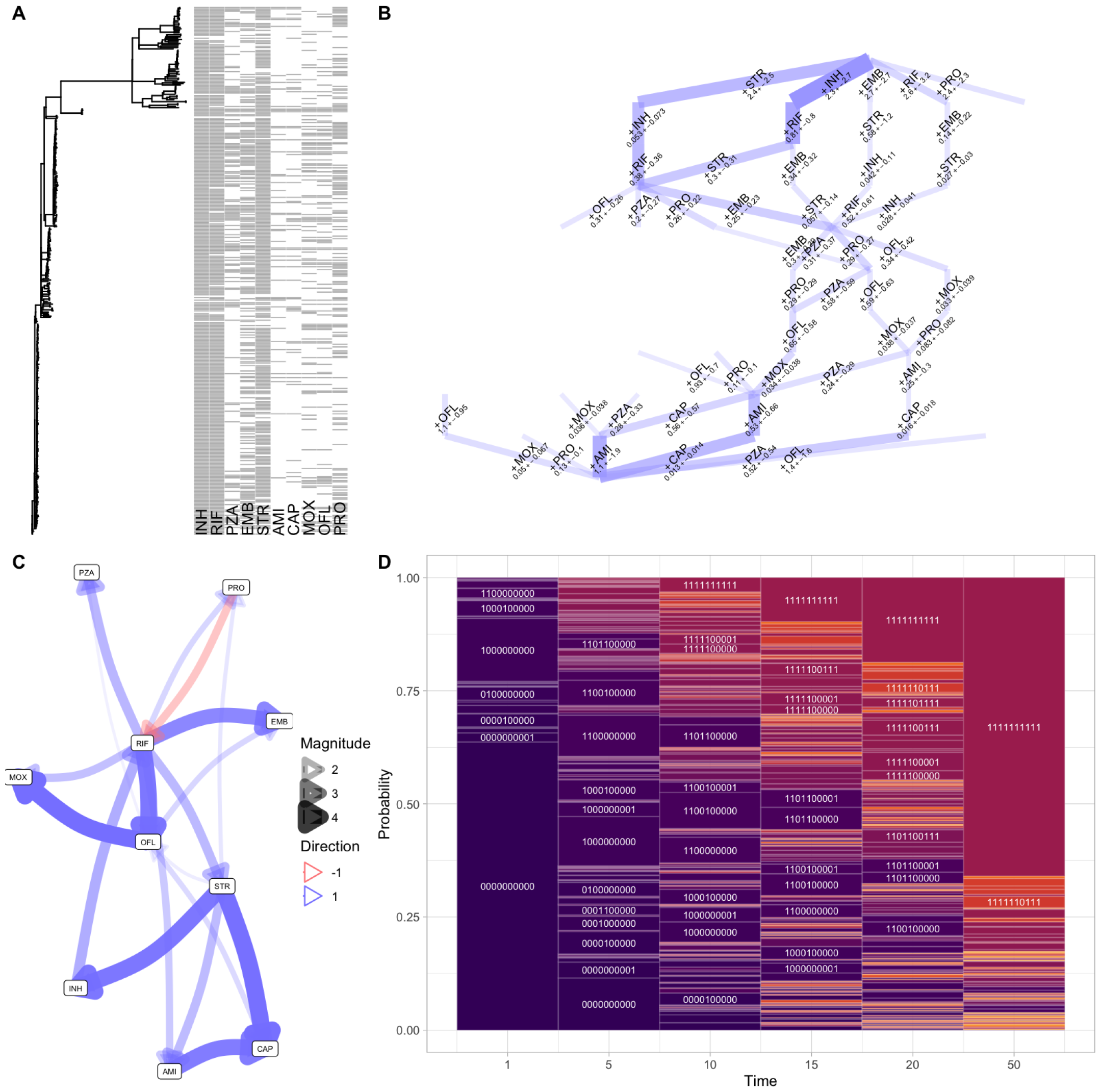
Inference of the dynamics of anti-microbial resistance evolution in tuberculosis. **(A)** Dataset from Casali et al. (2014), consisting of a phylogeny connecting observations of drug resistance presence/absence. **(B)** Inferred high-probability transitions between states in the TB AMR system. Transitions are labelled by the drug to which resistance is acquired in that step, and inferred likely mutational ‘timescale’. Edge width gives the probability of a transition; only transitions above a threshold probability of 0.05 are plotted. **(C)** Influence network plot from the *L*^2^ (mutual hazards) picture, describing inferred positive and negative influences between resistance acquisitions. Only influences with a posterior coefficient of variation under 0.3 are plotted. **(D)** Motif plot describing probabilities of different observations in the inferred evolutionary dynamics. Further outputs of the inference process are shown in Supp. Fig. S6A-C.

#### Dataset size

HyperTraPS-CT’s sampling approach means that it does not suffer a combinatorial explosion of paths that must be considered as *L* increases. Although performing adequate sampling to characterise large systems is still a challenge, it is not an insurmountable one for some examples of reasonable size. HyperTraPS has been used successfully for *>* 120 features (Peach et al., 2021; Johnston and Williams, 2016). Here, we asked whether the algorithm could recover dynamic structure with limited observations of large feature sets. To this end, we increased *L* in the double-pathway model system above, and took exactly one observation from each transition on the associated network. Fig. 3B shows that although convergence takes some time, the continuous-time case can readily identify dynamic patterns with 70 features with only 140 observations (Fig. 3B); other large *L* cases are shown in Fig. S4.

#### Maximum likelihood and Bayesian approaches

As HyperTraPS-CT is concerned with the likelihood estimation for a particular parameterisation, the surrounding inference scheme using this likelihood is flexible. Previous HyperTraPS work was based on a Bayesian picture through Markov chain Monte Carlo (MCMC) (Johnston and Williams, 2016) or auxiliary pseudo-marginal MCMC (APM MCMC) (Greenbury et al., 2020; Murray and Graham, 2016). We now include approaches for more straightforward likelihood maximisation, including simulated annealing and stochastic gradient descent. These approaches return a single point estimate for a high-likelihood parameterisation, but typically require substantially less processor time than the Bayesian exploration of parameter space (Supp. Fig. S3).

The Bayesian setting allows prior information to be included in the inference process. HyperTraPS-CT currently allows this prior information to be specified in the form of uniform distributions on parameter values, allowing the scale and sign of individual rates and feature interactions to be specified or constrained. A simple example is given in Fig. 3C, where a prior is used to constrain the basal rates for each feature in the accumulation process. This has the direct posterior influence of rebalancing the probabilities of the initial steps in the accumulation process, as well as shifting the posterior probabilities of later transitions.

#### No, uncertain, or precise timing observations

Representation of observation times as time windows (which can have infinite, finite, or zero width) allows different degrees of uncertainty in observation time to be captured. This is of particular use, for example, in phylogenetically embedded data, where the branch lengths connecting an ancestor to a descendant are uncertain; or in cross-sectional snapshot data, where the time since the ‘root’ state is typically unknown. The effects of increasing or decreasing the uncertainty of observation times in the example system are demonstrated in Supp. Figs. S2E-F and S5; this ability is exploited in the tuberculosis case study below.

#### Predictions of hidden features and future transitions

HyperTraPS supports prediction of unseen features (for example, given an inferred transition network, what is most likely to replace the ?s in 001101??0?) and future dynamics (for example, given an inferred transition network, what is most likely to happen next to the state 001101010, and how long will it take?). Simple illustrative examples are given in Fig. 3 D. These predictions may be useful in applied settings: for example, if a new bacterial strain has some but not all of its drug resistance phenotypes analysed, and clinicians require a prediction of which resistance will evolve next.

#### Visualisations

As part of the HyperTraPS-CT software implementation, we have included a suite of visualisation procedures allowing interpretation of inference outputs. These include diagnostic traces of values during the optimisation or MCMC process (Supp. Fig. S3); summaries of the inferred dynamics by probabilities of different event orderings (Figs. 3B, 4C, Supp. Figs. S2, S3) and probabilities of different states over time (Figs. 2D, 4D); full or truncated visualisations of the inferred transition graph with associated timings (Figs. 2A, 3A,C, 5B, 4A); and visual reports of predictions from the inferred model (Fig. 3D). Depending on the model structure chosen, matrices or graphs of influences between individual features (Figs. 2B, 5C, 4B) or sets of features (Fig. 3A) can also be produced.

Taken together, these examples demonstrate the capacity of HyperTraPS-CT to work with (i) different model structures (positive and negative pairwise/mutual hazard interactions, influence of larger sets of features on accumulation dynamics, arbitrary logic interactions and completely independent transitions between states); (ii) different data structures (cross-sectional, phylogenetic, longitudinal data with absent, precise, or uncertain observation timings); (iii) different inference approaches (maximum likelihood, Bayesian, different regularisations); (iv) large datasets of many dozen features.

### Cancer progression in acute myeloid leukemia

To demonstrate HyperTraPS’ capacity alongside state-of-the-art alternatives, we first look to the cancer progression field, where accumulation modelling is well established (Diaz-Uriarte and Herrera-Nieto, 2022). For a comparison with a recent approach, itself compared positively with previous approaches, we use the single-cell genomic dataset of acute myeloid leukemia evolution from (Morita et al., 2020), previously analysed with TreeMHN (Luo et al., 2023). This dataset consists of a set of trees linking ‘ancestral’ and ‘descendant’ observations, represented as barcodes describing the presence/absence of a mutation in a set of genes. Following ancestral state reconstruction on these trees, assuming a mutation-free precursor root state, we used HyperTraPS-CT with penalised likelihood to infer the pathways of mutation acquisition during cancer progression. Fig. 4A shows a truncated set of most probable early accumulation steps; Fig. 4B shows an inferred map of interactions between mutations; Fig. 4C shows the probability that a given mutation occurs at a given ordinal step through the accumulation process.

**Figure 4:**
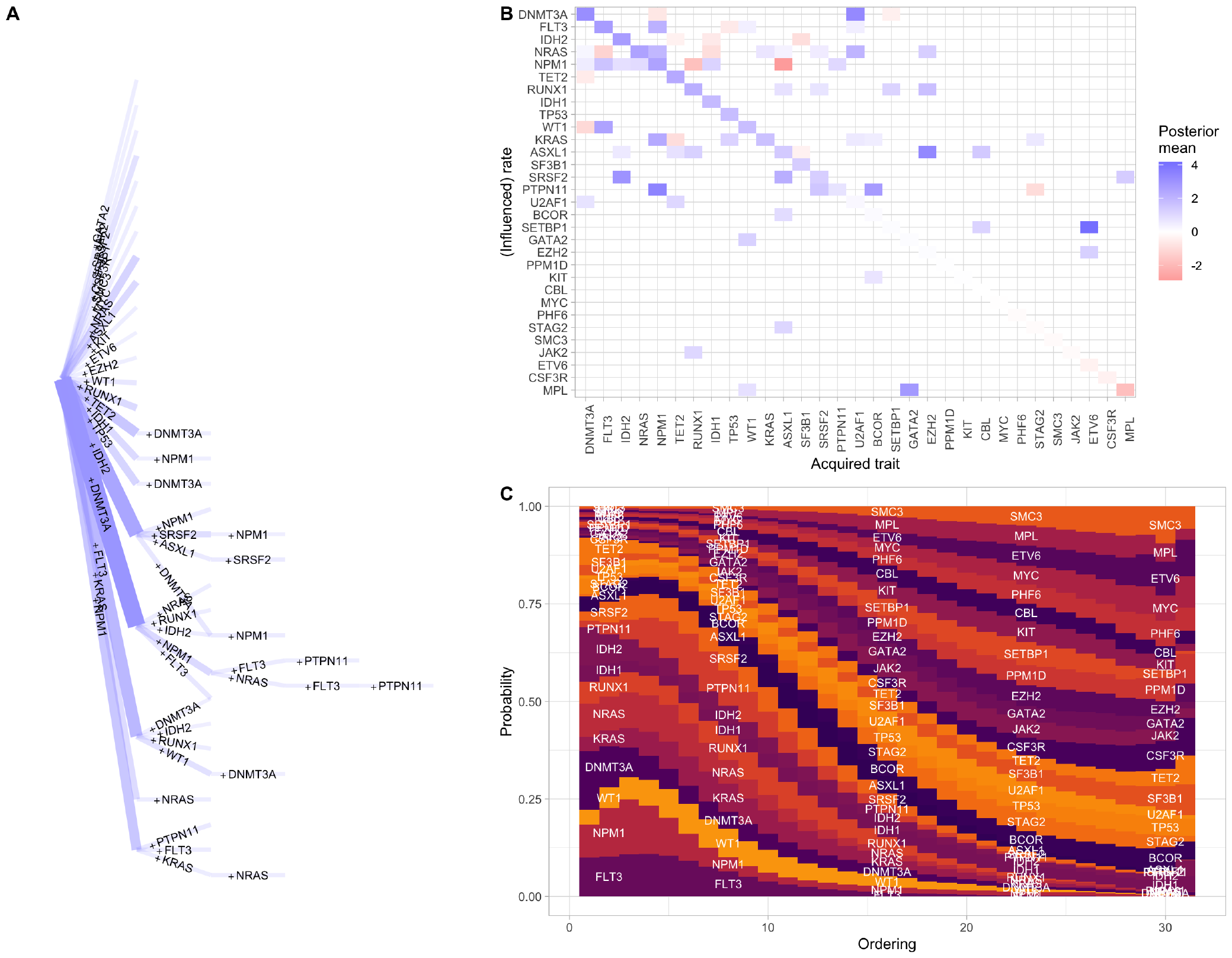
Accumulation of mutations driving cancer progression. **(A)** Truncated set of inferred high-probability transitions between states in the cancer progression system. Transitions are labelled by acquired feature; only transitions with a posterior probability over 0.006 are plotted. **(B)** Map of base acquisition rates and positive and negative pairwise influences between features (corresponding network plot in Supp. Fig. S7. **(C)** Motif plot showing the probability that a particular mutation is acquired at a given ordinal step in an accumulation process.

HyperTraPS-CT with penalised likelihood identifies many of the same features as TreeMHN. The ordering of base rates of mutational changes is very comparable: *DNMT3A* has the highest base rate, followed by *IDH2, FLT3, NRAS*, and *NPM1*, with the same collection of genes following these. Sets of interactions between changes are also consistently identified, with, for example, *DNMT3A* typically acting to promote the acquisition of other mutations (except *WT1*); *IDH2* having a more limited set of promotion partners; *FLT3* acting to both promote some mutations (*NPM1, WT1, RUNX1*) and suppress others (*NRAS*); many mutations acting to promote the acquisition of *NPM1* (except for suppressors *RUNX1* and *ASXL1*), and more features. The relative magnitudes of the inferred interactions are also largely consistent between the two approaches, with, for example, the influence of *FLT3* on *NRAS* being the strongest inferred suppression, and the influences of *IDH2* and *ASXL1* on *SRSF2* among the strongest inferred promotions.

The agreement with TreeMHN here (and behaviour in the simple case studies above) is positive support that HyperTraPS-CT provides a consistent way of estimating progressive accumulation dynamics with real as well as synthetic data. We next looked to a system where its ability to handle uncertain continuous timing data could be demonstrated.

### Acquisition of antimicrobial resistance genes in tuberculosis

To this end, we next used HyperTraPS-CT to explore an evolutionary question of pronounced global health importance – the evolution of antimicrobial resistance – in *Mycobacterium tuberculosis*, using a Russian study (Casali et al., 2014). We used the curated data used in Greenbury et al. (2020) where a set of bacterial isolates, related via a known phylogeny, have barcodes corresponding to the presence/absence of resistance to each of ten drugs (Fig. 5A). These drugs are referred to by three-letter codes: INH (isoniazid); RIF (rifampicin, rifampin in the United States); PZA (pyrazinamide); EMB (ethambutol); STR (streptomycin); AMI (amikacin); CAP (capreomycin); MOX (moxifloxacin); OFL (ofloxacin); and PRO (prothionamide). We used HyperTraPS-CT with penalised likelihood to learn the pathways of accumulating multi-drug resistance. We reconstructed ancestral states on the phylogeny using the maximum parsimony approach, assuming irreversible accumulation dynamics. The branch lengths on the phylogeny (Fig. 5A) do not correspond to absolute timings but to a measure of evolutionary change, estimated from independent genomic data (Casali et al., 2014; Greenbury et al., 2020). We first assume that this estimate phylogeny is precise, and use a continuous time inference picture precisely specifying observation time as the branch length *b* for each transition (*τ*_1_ = *τ*_2_ = *b* in Eqn. 1). The results are shown in Fig. 5.

Beginning with the inferred accumulation pathways on the hypercubic transition graph (Fig. 5B), resistance to INH, then RIF/STR in either order, then likely EMB is the dominant pathway. This highest-probability pathway matches observations from discrete-time inference in Greenbury et al. (2020) and Moen and Johnston (2023) (also shown in Supp. Fig. S6D). However, accounting for the continuous ‘time’ picture suggests another mode which is less prominent in the discrete-time picture: the early acquisition of STR resistance, followed by INH, then RIF. This pathway emerges in the continuous time picture because, although STR resistance is less common than INH or RIF resistance in the dataset, the transitions involving early STR resistance occur on a shorter timescale. This information is of course excluded from discrete-time pictures.

Following these early steps, pathways become more branched, with a collection of competing routes and corresponding timescales visible in Fig. 5B,D. Generally PRO and PZA resistances are likely acquired at intermediate stages, and MOX, OFL, CAP, AMI at later stages. The modal final resistance acquisition sequence is PZA-CAPAMI. The general ordering of these resistances agrees with Greenbury et al. (2020), but the relatively strong support for AMI (not CAP) as the final resistance step again emerges because of the timings of the associated observations. The map of interactions (Fig. 5C) gives a collection of positive interactions between features, and one negative interaction (PRO repressing RIF acquisition).

What if we cannot assume the phylogeny linking these observations is precisely known? If branch lengths are uncertain, the associated observation times for an ancestor-descendant transition are also subject to uncertainty. To demonstrate how HyperTraPS-CT can allow for such uncertainty, we set *τ*_1_ = 0, *τ*_2_ = *b* instead of *τ*_1_ = *τ*_2_ = *b* above. In this way, the observation of the descendant state at some time between 0 and *b* is required, rather than precisely at *b*. The corresponding outputs are shown in Supp. Fig. S6E-F. The same dynamics are recovered but with substantially increased uncertainty on transition timescales and fewer robustly supported interactions, reflecting the additional uncertainty in the system.

We also used the model flexibility in HyperTraPS-CT to explore evidence for feature interactions beyong the pairwise cases identified in Fig. 5. To this end, we used the *L*^3^ model (allowing nonadditive influence of feature pairs on other feature acquisitions) with penalised likelihood on the same tuberculosis dataset. Supp. Fig. S6G shows that some nonadditive influences of feature pairs on acquisition rates were identified with confidence: for example, a combination of RIF and OFL positively influences the rate of acquisition of MOX.

### Prediction of unobserved and future behaviours in accumulation dynamics

The inferred dynamics here can be used, for example, to provide predictions about the likely next drug resistance to evolve from a given observed state (as in Fig. 3Dii), and to predict phenotypic features from an imperfectly sampled new observation (as in Fig. 3Di). Both of these predictive statements could be of conceivable applied use. A prediction that resistance to a given drug is likely to occur next might suggest the use of a different drug (one with lower and/or later acquisition propensity). In cases where phenotypic assays of drug resistance are challenging (requiring researcher time and resources), the predictions of unmeasured drug resistances could be used as a (clearly imperfect) substitute.

To test the capacity of HyperTraPS-CT to make such predictions, we split the transition set for the tuberculosis case study into 75:25 training:test subsets, and used the training subset to produce posteriors on the hypercubic model. We did this 75:25 split ten independent times to guard against accidents of sampling: the following results are amalgamated across these instances. To address the first predictive task, using the unseen test data, we identified transitions where the ‘ancestral’ and ‘descendant’ states differed by one acquired feature, and asked whether acquisition of that inferred feature was among the most probable predicted steps from the ancestral state. In the majority of cases, the true acquisition was the most or next-most likely step predicted by the trained model, with the most-likely predicted step being clearly most common (Fig. 6A).

**Figure 6:**
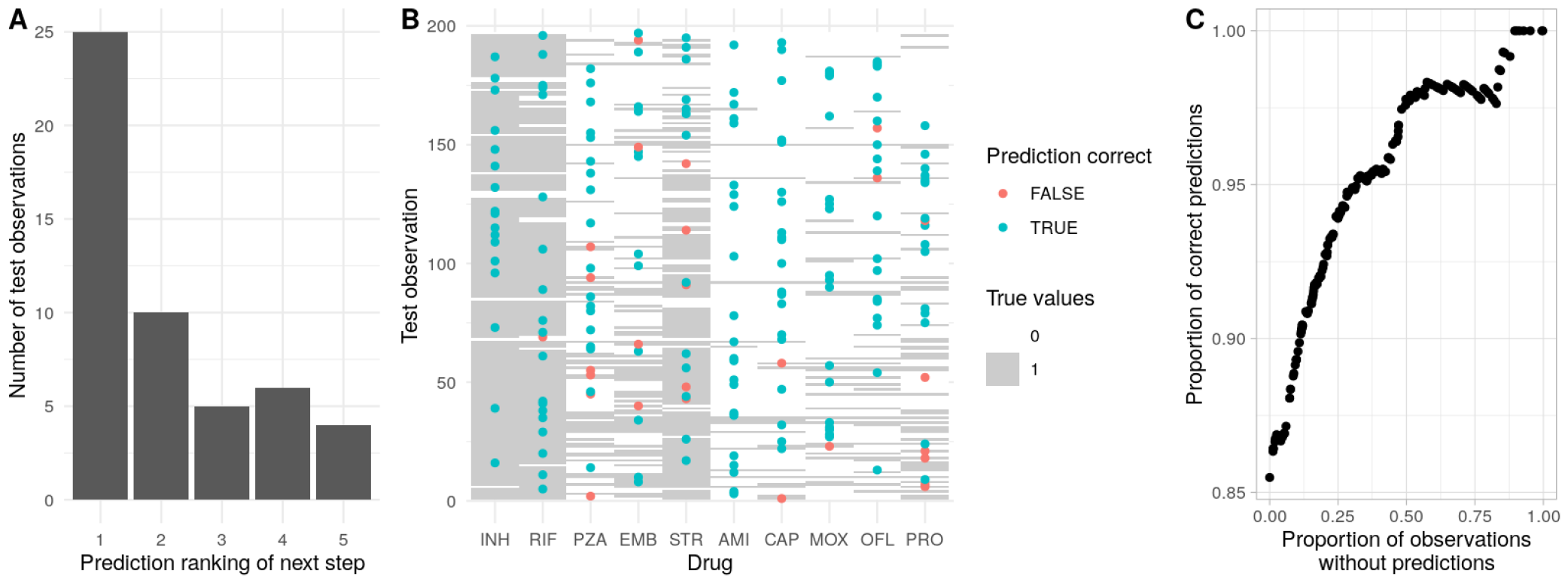
Predictions of future and unobserved features in tuberculosis multi-drug resistance evolution. **(A)** Following training, predictions were made for the next accumulation step in a set of withheld test observations (see text). For each observation, each possible step from the precursor state was ranked by predicted probability. The plot shows the predicted ranking of the true acquisition (1 predicted to be most likely) across test observations. **(B)** Prediction of artificially obscured features in test data. The ‘barcode’ background shows the full test data, the points correspond to an artificially obscured feature that was predicted from the trained model, coloured by whether the prediction was correct or not. 86% of predictions were correct in this example. **(C)** More generally, a parameterised choice can be made about whether to assign a confident prediction based on its posterior probability. This plot shows how the proportion of correct predictions increases as this ‘strictness’ criterion is increased, at the cost of a higher proportion of observations without an assigned prediction.

We next artificially obscured a random set of features in the test data and attempted to predict their values based on the remaining features and the model inferred from the training data (Fig. 6B). For each obscured feature we recorded *P* (1), the posterior probability that it would take value 1, from the inferred model. With a ‘strictness’ parameter Δ, we assigned a prediction of 1 for features with *P* (1) *>* 0.5 + Δ, a prediction of 0 for *P* (1) *<* 0.5 − Δ, and no prediction otherwise. As Δ changes from 0 to 0.5 we thus require stronger posterior evidence for making a prediction. Fig. 6B shows the case for Δ = 0, where 86% of predictions were correct; the general behaviour of prediction accuracy against proportion of assigned predictions is shown in Fig. 6C. Enforcing stricter criteria for assigning a prediction readily amplifies accuracy over 95% at the cost of roughly a quarter fewer confident predictions.

Taken together, HyperTraPS-CT has demonstrated both compatibility with existing approaches for the study of cancer progression and in synthetic case studies, and the ability to exploit timing information to refine estimates of evolutionary dynamics in the study of anti-microbial resistance. At the same time, its flexibility will allow a range of scientific questions to be explored across other applied fields. These include the role of higher-order interactions between features in determining accumulation behaviour, and (with approaches to compare inferred transition graphs (Garcia Pascual et al., 2024)) the similarities, differences, and modulating factors of accumulation dynamics in different samples.

## Discussion

We have introduced a continuous timescale to hypercubic transition path sampling (HyperTraPS), a flexible approach to inferring the dynamics of accumulaton models. We have also generalised the parameter structures used in the underlying model so that arbitrarily high orders of interaction between feature sets can be captured by the inference process, and introduced a panel of new options for predictions, inclusion of prior knowledge, optimisation approaches, regularisation, and visualisation. Higher-order interactions have previously been shown to have explanatory power in, for example, ovarian cancer progression (Moen and Johnston, 2023), and limited higher-order interactions between drug resistances are observed here in the tuberculosis case study. We hope that the regularisation and model selection approaches we provide here will help explore potentially useful model structures in other contexts.

We have shown that the consideration of continuous timing information can lead to differences in the outcome of inference – as demonstrated by the ordering of streptomycin (STR) resistance in the tuberculosis case study. Features that are less represented in a dataset, but which are associated with rapid acquisition times when they do appear, will tend to be assigned later acquisition orderings in explicitly or implicitly discrete-time approaches. Timing information can help resolve these dynamics – but is typically uncertain, as feature acquisitions can occur anywhere between two sampling events. We have outlined and demonstrated one approach by which such uncertainty may be addressed – through including a time window of controlled length during which an observation may be made. This approach corresponds to a particular error model – a uniform distribution over possible observation times. Generalising this model to include different distributions over observation times constitutes a target for future work. The robustness of our approach can also be tested by artificially perturbing the source dataset and assessing the influence of these perturbations on the posteriors (Johnston and Williams, 2016).

As with other approaches, the posteriors that HyperTraPS initially produces reflect an ‘umbrella’ picture of accumulation dynamics, combining evidence from observations that may have been subjected to different selective pressures and environmental conditions (Greenbury et al., 2020; Johnston and Williams, 2016). This umbrella picture reflects coarse-grained posterior knowledge of accumulation pathways given all observations – in a sense, describing the distribution of belief about a putative new sample about which we have no further information. Refinement of these umbrella posteriors to a finer-grained picture is possible by including information about the context of different samples. Previous examples include the broader taxonomy of an individual in evolutionary dynamics (Williams et al., 2013), level of patient risk in disease progression dynamics (Johnston et al., 2019), and an individual’s ecology in behavioural dynamics (Johnston and Røyrvik, 2020). In such cases, observed multimodality in the umbrella posteriors can sometimes be accounted for (at least partly) by a natural distinction between data subsets from differently categorised observation, informing about mechanistic influences on the accumulation process.

In some contexts, the properties of some observed states may not be completely known. Uncertainty in descendant observations (and hence anywhere in cross-sectional experimental design) does not challenge our approach, which readily supports a ‘missing data’ flag in any number of features for such observations. However, uncertainty in ancestral states in the longitudinal or phylogenetic context poses a larger technical challenge (Johnston and Williams, 2016). Our approach can always be applied using a method originally called ‘phenotypic landscape inference’: uniform, unbiased sampling with likelihoods estimated by tracking the number of states compatible with uncertain observations (essentially using brute force sampling to estimate the left-hand side of Eqn. 1), as originally implemented in Williams et al. (2013). However, the question of how to *most efficiently* sample evolutionary paths from an uncertain origins will be addressed in future research.

HyperTraPS by its nature is a sampling approach, and does not have the native capacity for exact calculations of transition path probabilities as found in other approaches including MHN (Schill et al., 2020) and HyperHMM (Moen and Johnston, 2023). For small systems, the likelihoods estimated from this sampling approach are effectively indistinguishable from the exact results (Supp. Fig. S2), and the precision of this estimation can be controlled (at the expense of computational time) by the number of sampled paths *N*_*h*_. The inevitable expansion of pathway and parameter space as the number of features *L* increases means that sampling approaches are currently used even when using MHN for large systems (Luo et al., 2023). However, the necessarily random nature of the sampling underlying HyperTraPS must be considered in its responsible interpretation; convergence of results from different random number seeds, for example, should be assessed as a ‘safety check’. On a related note, cases will clearly exist in accumulation modelling where different parameterisations may have equal abilities to reproduce observations. The Bayesian embedding of HyperTraPS, and the approaches for regularisation and model comparison we suggest here, can be used in such cases where identifiability is challenged, to allow reporting of the flexibility and constraints on different mechanistic parameters under different model structures.

As approaches for accumulation modelling, traditionally grounded in cancer statistics, gain more and more features in common with evolutionary biology methods (including connections with trees/phylogenies, and continuous time), it is worth reiterating that methods for discrete Markov dynamics on trees have existed in the evolutionary literature for decades (Pagel, 1994; Lewis, 2001; Harmon, 2019). The Mk (Markov k-state) model is well established in evolutionary biology and any discrete Markovian model on a phylogeny (including a ‘star’ phylogeny, corresponding to independent instances / cross-sectional observations) can in principle be viewed as a subset of this picture (Boyko and Beaulieu, 2021). In accumulation modelling the focus is typically on features and their relationships, rather than states and their relationships, and the different dependency structures (pairwise interactions, logic interactions, and so on) are often explicitly encoded in accumulation models in a way that would be less straightforward to extract – and potentially impossible to infer in reasonable computational time – from an Mk model picture. But the connections between the disciplines will be worth exploring in future developments.

## Acknowledgments

The authors are grateful to Ellen Røyrvik, Paul Una, Amelia Earl, and Ai Monti for valuable discussions. This project has received funding from the European Research Council (ERC) under the European Union’s Horizon 2020 research and innovation programme (Grant agreement No. 805046 (EvoConBiO) to IGJ). This work was supported by the Trond Mohn Foundation [project HyperEvol under grant agreement No. TMS2021TMT09 to IGJ], through the Centre for Antimicrobial Resistance in Western Norway (CAMRIA) [TMS2020TMT11]. RDU was partially supported by grant PID2019-111256RB-I00 funded by MCIN/AEI/10.13039/501100011033.

## Supplementary Information

### HyperTraPS in continuous time

To extend HyperTraPS to analyse continuous timing, we first consider the targets of inference and structure of data that we will be dealing with. The state space *H* of the system is of size 2^*L*^ and each state *s* is labelled by a unique binary string of length *L*. Let *λ* be a transition matrix where *λ*_*i→j*_ ≡ *λ*_*ij*_ is the intensity of a transition from state *i* to state *j*. All transitions are assumed to be Poissonian. Define a path *c* in *H* as a sequence of *n* states *c*_1_, …, *c*_*n*_.

If we work in a picture of feature accumulation, dynamics can be envisaged as originating from an ‘ancestral’ state possessing no features (000… ≡ 0^*L*^). Then *λ*_*i→j*_ is nonzero only for pairs where *j* differs from *i* by exactly one 0 → 1 change. In an alternative picture of feature losses, the ancestral state possesses all features (111… ≡ 1^*L*^) and *λ*_*i→j*_ is nonzero only for pairs where *j* differs from *i* by exactly one 1 → 0 change. The two pictures can be naturally linked by considering a loss as an ‘acquisition of absence’.

To build a framework where both the original and continuous-time cases can be analysed together, we consider as the fundamental quantity of interest *P* (*b, τ*_1_, *τ*_2_| _*a*_, 0; *λ*) – *the probability that, having started at state a at time* 0, *the system will be in state b at a time between τ*_1_ *and τ*_2_, *given a transition matrix λ*. When *τ*_1_ = 0, *τ*_2_ = ∞, this gives the original HyperTraPS expression for *P* (*a* → *b*); in the limit *τ*_2_ → *τ*_1_, this requires that the system is in state *b* at the particular time *τ*_1_. From here on, for clarity, we will treat the *λ* dependence as implicit.

HyperTraPS gives the probability that, having started at *a*, the system will encounter *b* (hence, *P* (*b*, 0, ∞| _*a*_, 0)). To compute *P* (*b, τ*_1_, *τ*_2_| _*a*_, 0) in more general cases we need additional probabilities, specifically, (i) the probability that we arrive at *b* before or at *τ*_1_, and do not then move from *b* until at least *τ*_1_, and (ii) the probability that we arrive at *b* between *τ*_1_ and *τ*_2_:

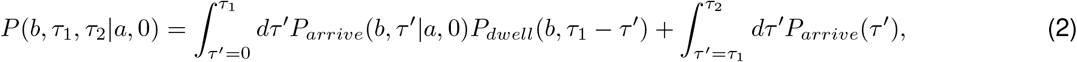

where *P*_*arrive*_ and *P*_*dwell*_ respectively give arrival and ‘dwell time’ probabilities: that is, the probability that *b* is first reached at time *τ* ^*′*^, then remains there for at least a further *τ*_1_ − *τ* ^*′*^ period.

It will be useful to define the characteristic ‘escape’ rate *β*_*i*_ for leaving a point *i* in the transition network, which is simply the sum of rates of processes leaving that point, hence

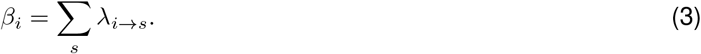

We first consider *P*_*dwell*_(*b, τ*), the probability that we dwell at *b* for at least time *τ*, which is straightforward to compute. As all the processes by which we can leave *b* are Poissonian and independent, the process ‘leave *b* by any method’ is also Poissonian, with rate *β*_*b*_. Then the dwell time is exponentially distributed:

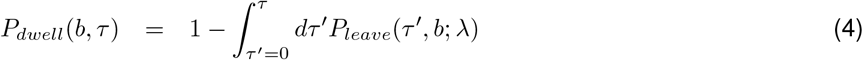

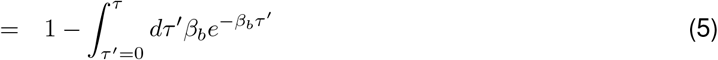

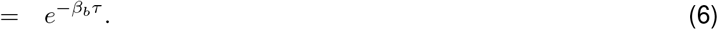

We now turn to *P*_*arrival*_. Consider a particular path *c* across the transition network, where steps are Poisson processes with rates 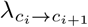, the first state *c*_1_ ≡ *a*, and the *n*th step reaches a target *c*_*n*_ ≡ *b*. The probability distribution of arrival times at *b* is then the distribution of the sum of *n* independent exponentially-distributed random variables describing the waiting times for each step, which is a hypoexponential distribution (Ross, 2014). For a general set of states:

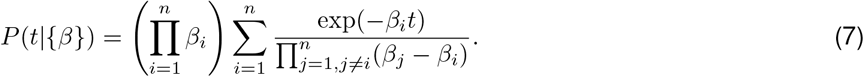

Hence, the probability that a path *c* reaches its final point *c*_*n*_ at time *t* is

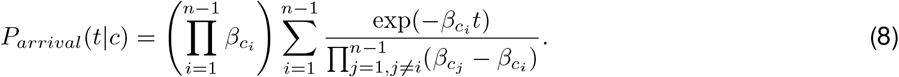

For clarity we define

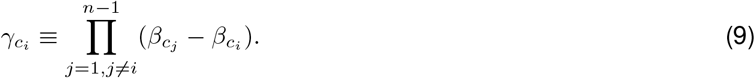

It will be useful later to consider

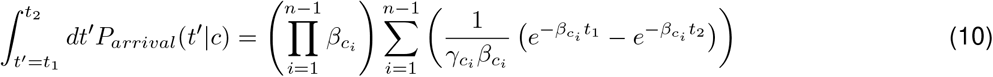

It is straightforward to verify that setting *t*_1_ = 0, *t*_2_ = ∞ gives 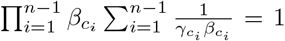; thus, the probability that the final step is reached at some finite time is 1 unless any 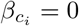.

The overall arrival time distribution at *b* is then given by a weighted sum of this distribution for each path that does indeed arrive at *b*, hence

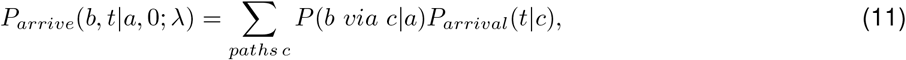

where *P* (*b via c*|*a*) is to be determined, but is clearly zero if path *c* does not lead from *a* to *b*.

The generally large number of possible paths on the transition network that do not lead from *a* to *b* will lead to this expression being hard to sample naively. Following HyperTraPS, we therefore consider how to make progress sampling only those paths of interest.

Define a state *s* to be compatible with a state *b* if, for a picture of evolutionary losses, there exists no *i* for which *s*_*i*_ = 0 and *b*_*i*_ = 1 (for the alternative picture of evolutionary acquisitions, this inverts to require no *i* for which *s*_*i*_ = 1 and *b*_*i*_ = 0). Thus, *s* is compatible with *b* iff *b* can be reached from *s* on the hypercube digraph. Define by *B*(*s*) the set of states accessible by one step from state *s* that are compatible with *b*. HyperTraPS gives us the probability of a path that is guaranteed to encounter *b* as

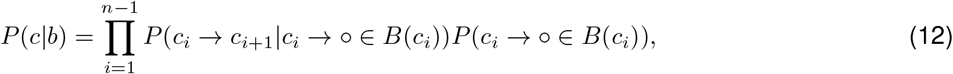

where º denotes ‘any element’, and a sampling scheme that encounters only paths leading from *a* to *b*, with probability

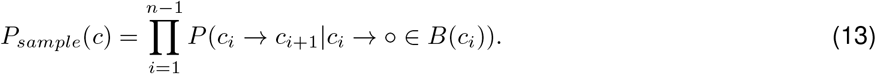

Recording Π_*i*_ *P* (*c*_*i*_ → º ∈ *B*(*c*_*i*_)) through a path sampled by HyperTraPS thus gives an estimate for *P* (*c*|*b*) for that path, which is guaranteed to lead from *a* to *b*, and thus gives *P* (*b via c*|*a*). We can then record *P*_*arrival*_(*t*|*a* → *b via c*) for HyperTraPS-sampled paths *c* and weight each according to its *P* (*c*), computed with Eqn. 12:

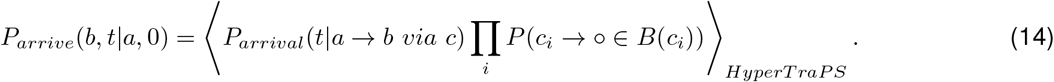

**Figure S1:**
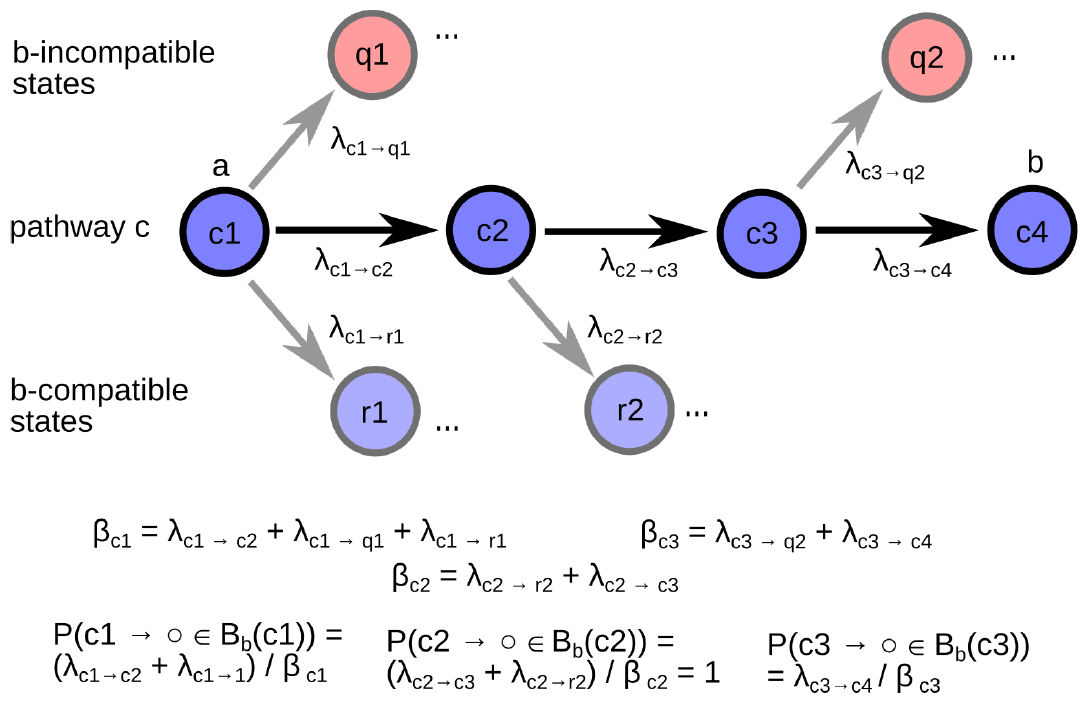
Computing *β* and *α* for a pathway *c*. An illustrative pathway running from *a* ≡ *c*_1_ to *b* ≡ *c*_4_. At each step, alternative transitions may exist to *b*-compatible states (*r*_1_, *r*_2_) and/or *b*-incompatible states (*q*_1_, *q*_2_). The escape rate *β*_*s*_ for a given step on the pathways is the sum of rates leaving that state; the compatibility probability *P* (*s* → º ∈ *B*_*b*_(*s*)) is the probability of undergoing a transition to a *b*-compatible state. The compatibility product *α* is the product of this probability over every step in the pathway. *α* and the set of *β* values are the statistics of a pathway that are used in constructing the estimated transition probability (Eqn. 1).

Consider replacing *P*_*arrival*_(*t*|*a* → *b via c*) by a simple indicator function *I*(*c*) reporting whether path *c* hits *b*. The quantity reported will then be the probability that *b* is reached at any time: ∫_t*′*_ *dt*^*′*^*P*_*arrive*_(*b, t*^*′*^|*a*, 0), or simply *P* (*a* → *b*). All paths sampled by HyperTraPS will pick up a unit coefficient from the indicator function, and the original HyperTraPS expression for pathway probability is recovered.

We are now in a position to construct the expression required in Eqn. 2:

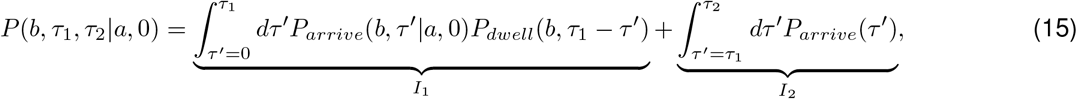

where the first integral *I*_1_ accounts for the system arriving at *b* before *τ*_1_ and dwelling there until *τ*_1_, and the second integral *I*_2_ accounts for the system arriving at *b* between *τ*_1_ and *τ*_2_.

The first integral is given by

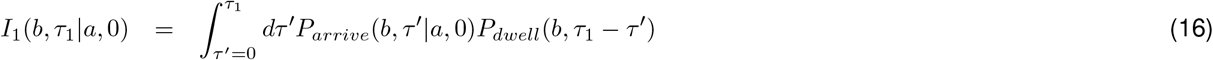

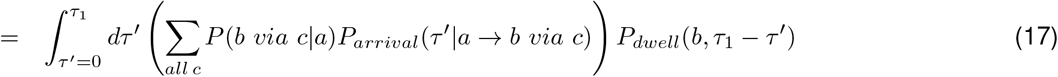

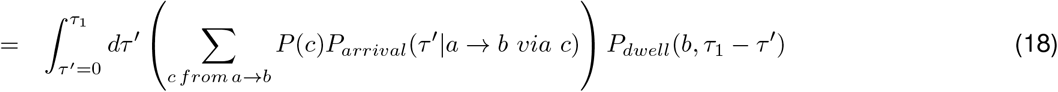

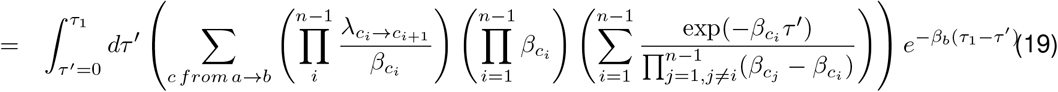

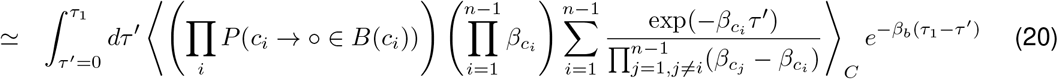

where the angle brackets denote sampling from a set of paths *C* using HyperTraPS. Noting that ⟨*f* (*c*)⟩_*C*_ = *f* (*c*)*N* (*c*)*/N*_*h*_, where *N* (*c*) is the number of occurrences of path *c* in *C* and *N*_*h*_ is the total number of HyperTraPS samples in *C*, this integral can be evaluated to give an estimate for *P* (*b, τ*_1_, *τ*_2_|*a*, 0):

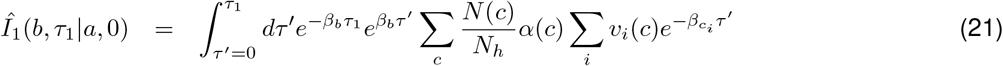

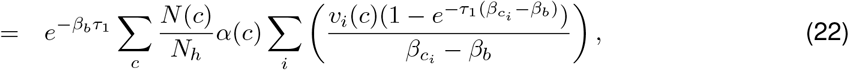

Where

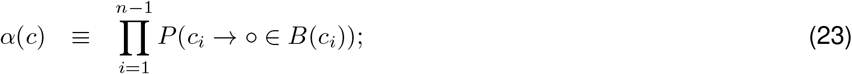

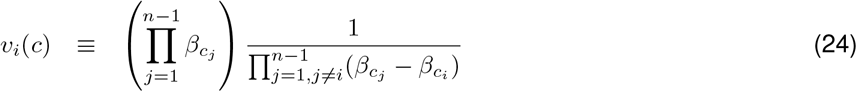

for *c*, a particular path sampled by HyperTraPS, and where the required *α* and 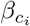 values can be easily recorded during a sampling algorithm.

If *a* ≡ *b*, the situation reduces to the case where we have arrived at *b* at *t* = 0, equivalent to *P*_*arrive*_(*b, τ* ^*′*^|*a*, 0) = *δ*(*τ* ^*′*^) in Eqn. 16. We are then only concerned with dwelling at *b* for the remaining time *t*:

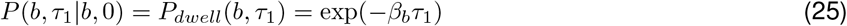

The second integral, *I*_2_, is given by

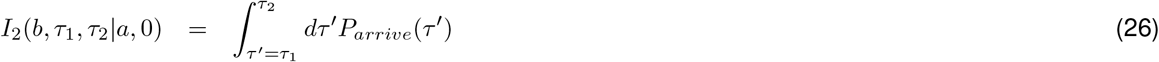

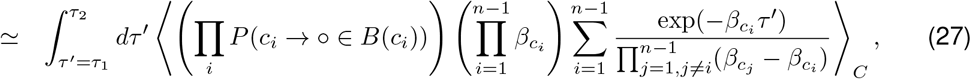

by analogy with the above derivation. Using Eqn. 10, this integral can also be solved to give

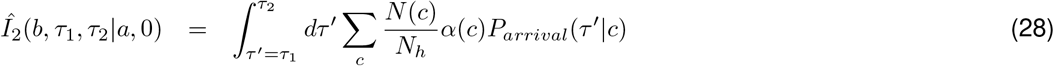

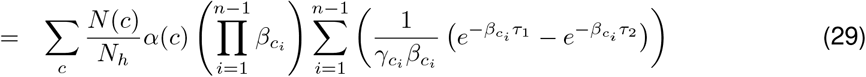

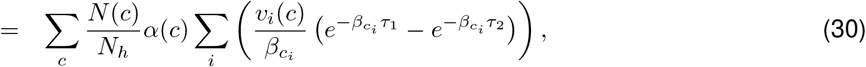

where, as before, the required *α* and 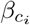 values can be easily recorded during a sampling algorithm. Together we finally obtain

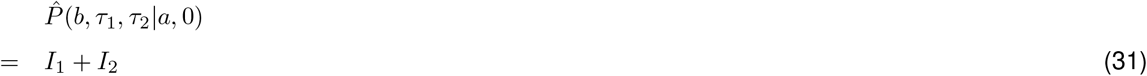

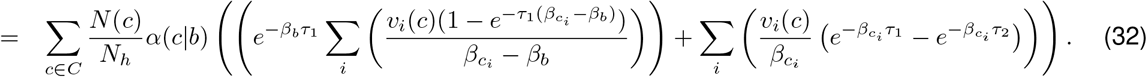

If complete sampling of the set of paths from *a* to *b* is possible, we can obtain an exact expression for *P* (*b via c*|*a*) in Eqn. 11. This probability is simply *P* (*c*)*I*(*c*|*a, b*), where *P* (*c*) = Π_*i*_ *P* (*c*_*i*_ → *c*_*i*+1_) and *I*(*c*|*a, b*) returns 1 if path *c* starts at *a* and ends at *b* and 0 otherwise. Following Eqn. 11, this expression can be substituted in Eqn. 24 for the approximate sampling weight *N* (*c*)*/N*_*h*_, giving

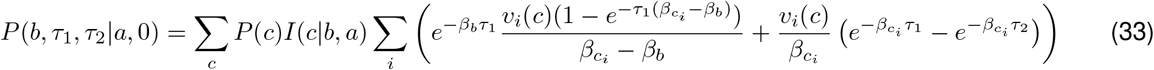

We can then run the HyperTraPS sampling algorithm to perform inference (i) in the absence of temporal information (*τ*_1_ = 0, *τ*_2_ = ∞, *I*_1_ = 0, *I*_2_ = *P* (*a* → *b*)); (ii) for precisely defined temporal samples (*τ*_1_ = *τ*_2_ = *τ, I*_1_ = *P* (*b, τ* |*a*, 0), *I*_2_ = 0); and (iii) for uncertain temporal samples (0 ≤ *τ*_1_ *< τ*_2_ *<* ∞, *I*_1_, *I*_2_ nonzero). Case (i) corresponds to original HyperTraPS. We note that the posterior orderings obtained from (i), (ii), and (iii) may in general differ.

## Inference using HyperTraPS-CT likelihood estimation

### Bayesian or maximum likelihood inference

Previous implementations of (discrete-ordering) HyperTraPS used a Bayesian approach with MCMC to identify transition matrices (given uniformative prior distributions) compatible with observations (Greenbury et al., 2020; Johnston and Williams, 2016; Williams et al., 2013). This process can naturally be extended to the continuous time picture (Fig. 1C).

#### Algorithm 2. Inference of posterior transition matrices using HyperTraPS-CT. Requires data D, prior distribution *P*_*prior*_(*λ*), MCMC parameters *θ*.

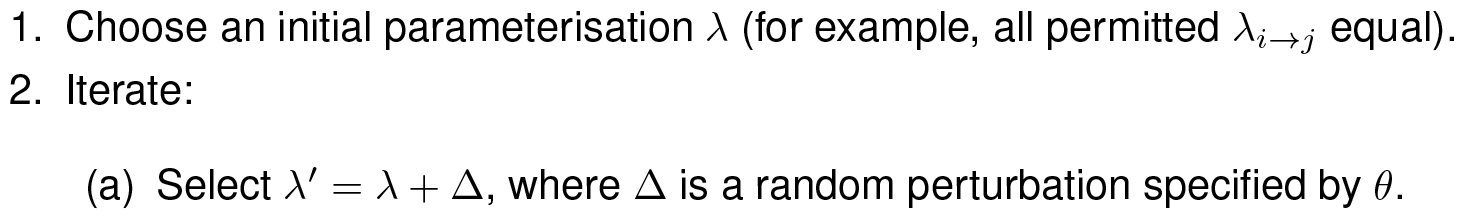
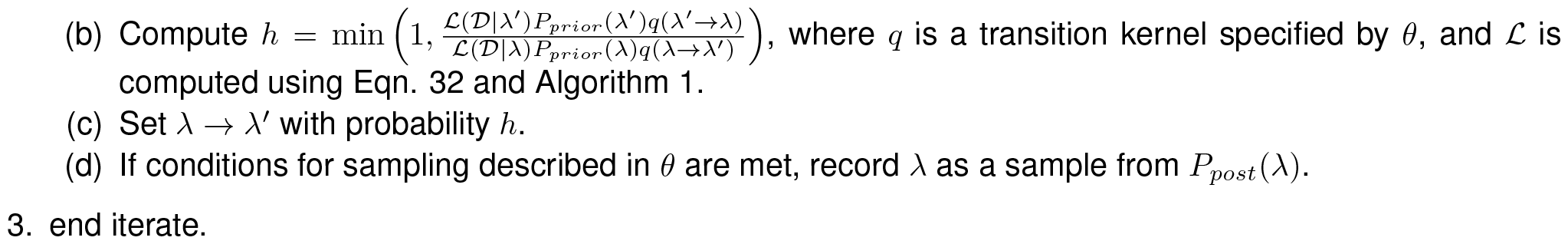

We note that in practise it is usually more convenient to work with log-likelihoods *ℓ* = log L than with L. We typically use uninformative uniform priors over log *λ*_*ij*_, and allow the parameters of the process above to be set by the user to best support converged inference results for a given dataset. A rule-of-thumb first choice is a perturbation kernel 𝒩 (0, 0.05) for log *λ*_*ij*_, *N*_*h*_ = 200 and 10^4^ MCMC iterations (discarding the first 20% as burn-in). If a maximum likelihood perspective is more desirable for reasons of computational time and/or interpretation, the likelihood estimate from HyperTraPS-CT can readily be maximised over parameters using any numerical optimiser. We include simulated annealing and a variant of stochastic gradient descent in the HyperTraPS-CT code; the behaviour of these approaches for the double-pathway test case is shown in Fig. S3.

### Parameter spaces

As *L* increases, the dimensionality of *λ* increases considerably. For tractability and/or interpretation, it may desirable to impose relationships between the different edges on the hypercubic transition network, and so decrease the dimensionality of the parameter space involved.

We begin by considering the dimensionality-reduction approach in Johnston and Jones (2016) and Hjelm et al. (2006), independently published as mutual hazard networks (Schill et al., 2020). Here, an *L* × *L* matrix is used to encode intensities for individual feature losses and the influence of one feature’s presence or absence on the loss intensity of another. Higher-order effects are neglected, and we no longer have an independent and unconstrained *λ*_*i→j*_ intensity for each transition, but the scaling of parameter space size with *L* is reduced from exponential to polynomial and subtle dynamics can still be captured (see below and Johnston and Williams (2016)). Specifically, if *s*^(*i*)^ is the binary absence/presence value of the *i*th trait in state *s*,

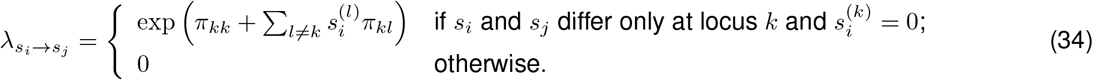

Hence, the diagonal elements *π*_*kk*_ encode a basal rate with which trait *k* is acquired, and the off-diagonal elements *π*_*kl*_ encode how the presence of the *l*th trait influences (positively or negatively) the rate with which trait *k* is acquired.

For example, *π*_11_ = 0, *π*_*ij*_ = Δ if *j* − 1 = 1, *π*_*ij*_ = −Δ otherwise, encodes a transition matrix with strong support for a single pathway, where trait 1 is acquired first (rate *π*_11_ = *e*^0^ against *e*^*−*Δ^ for other traits). The rate of acquisition of trait 2 then increases to *π*_22_ + *π*_12_ = 0, making trait 2 the next most likely acquisition, and so on.

For the next case, we consider a 3rd-order array *π*_*ijk*_. Then the ‘base rate’ for *k* is *π*_*kkk*_, the influence of *j* on *k* is *π*_*jjk*_, and the independent influence of the pair (*i, j*) on *k* is *π*_*ijk*_ = *π*_*jik*_. Clearly in this, and higher-order, cases, it is not true that every element of the corresponding array is independent: symmetries amongst the definitions of subsets (pair (*i, j*) is identical to pair (*j, i*)) mean that some elements have the same meaning. But in general we can use a subset of elements in the *n*th-order array to encode influences from single features, pairs of features, triples of features, and so on, as well as their ‘target’ base acquisition rate.

For the limiting case where transitions between states are completely independent – corresponding, in the feature picture, to arbitrary influences of subsets of all sizes – we simply use an array of size *L*2^*L−*1^, the cardinality of the edge set of the *L*-hypercube, to store the individual rates.

**Figure S2:**
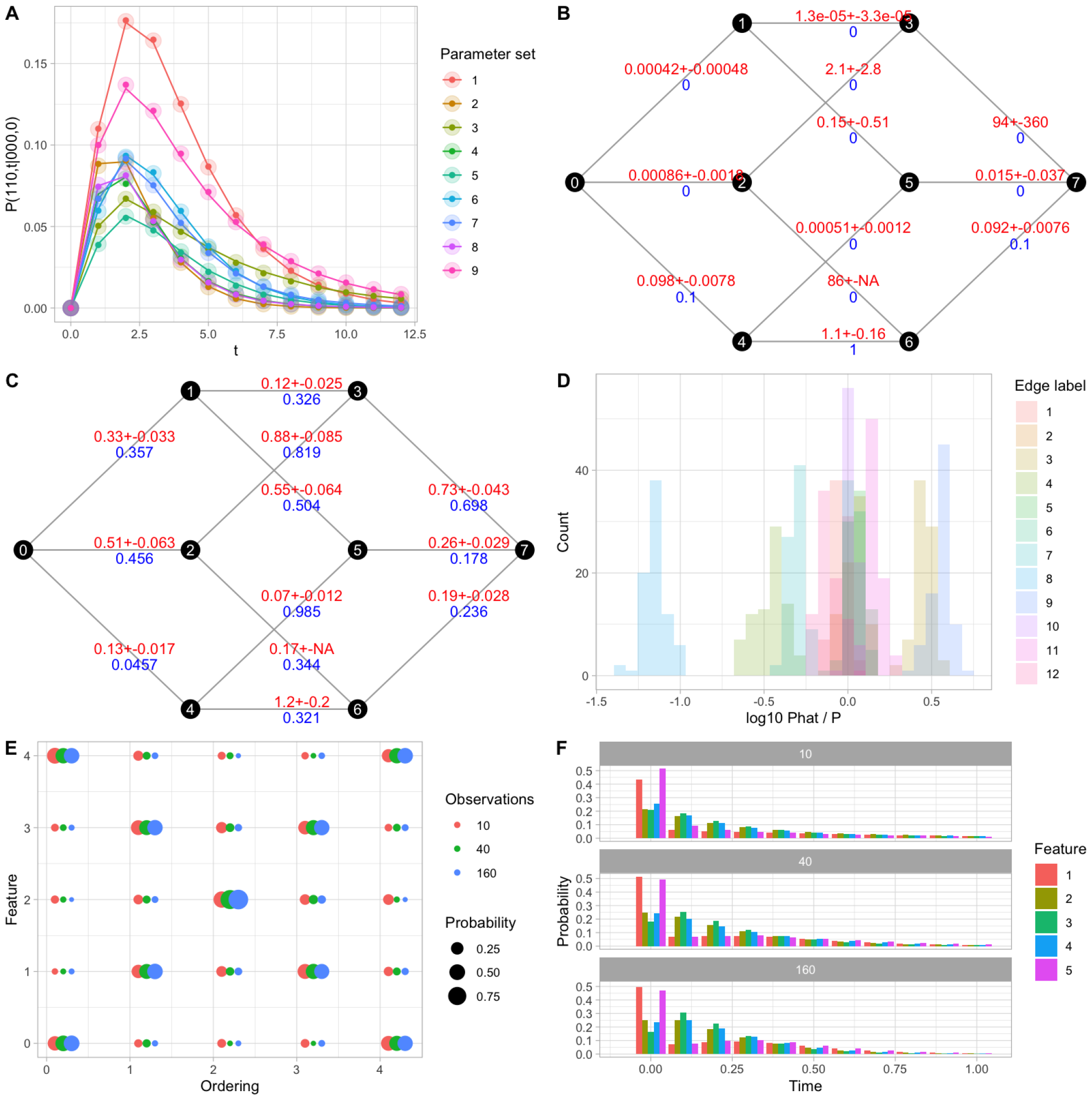
Validation and demonstration of HyperTraPS-CT (see text in ‘Validation, inclusion of prior information, and analysis of posteriors’ below). (A) Estimation of transition timescales and probabilities using HyperTraPS-CT (Eqn. 1; Algorithm 1, solid points) matches results from analytic calculation (lines) and exhaustive sampling (transparent points) of a tractable *L* = 3 model system under many different random parameterisations (each trace corresponds to a different parameterisation). (B) HyperTraPS-CT accurately recovers the original transition rate parameters for a given *L* = 3 system. Here, states are represented by their decimal equivalent (0 is 000, 6 is 110). Blue figures give the true system; red shows inferred means and 95% credibility intervals for each transition. (C) HyperTraPS-CT recovers original transition rates for a more complex model system; plots show original and inferred rates as in (B). (D) Inferred distributions of transition weights from (C). Histograms show the posterior distributions on each transition rate normalised by its true value; most distributions (blue) have modes in a fold range around the true value. (E) HyperTraPS-CT recovers posterior orderings of events for a model *L* = 5 system constructed to allow two competing paths (corresponding to an X-shaped structure in this plot). Smaller sample sizes (*n* = 10) lead to increased uncertainty in the pathway structures; higher observation numbers readily identify pathways. (F) Inferred timing distributions of feature acquisitions in the experiments in (E); in the generating process, each step takes 0.1 time units.

**Figure S3:**
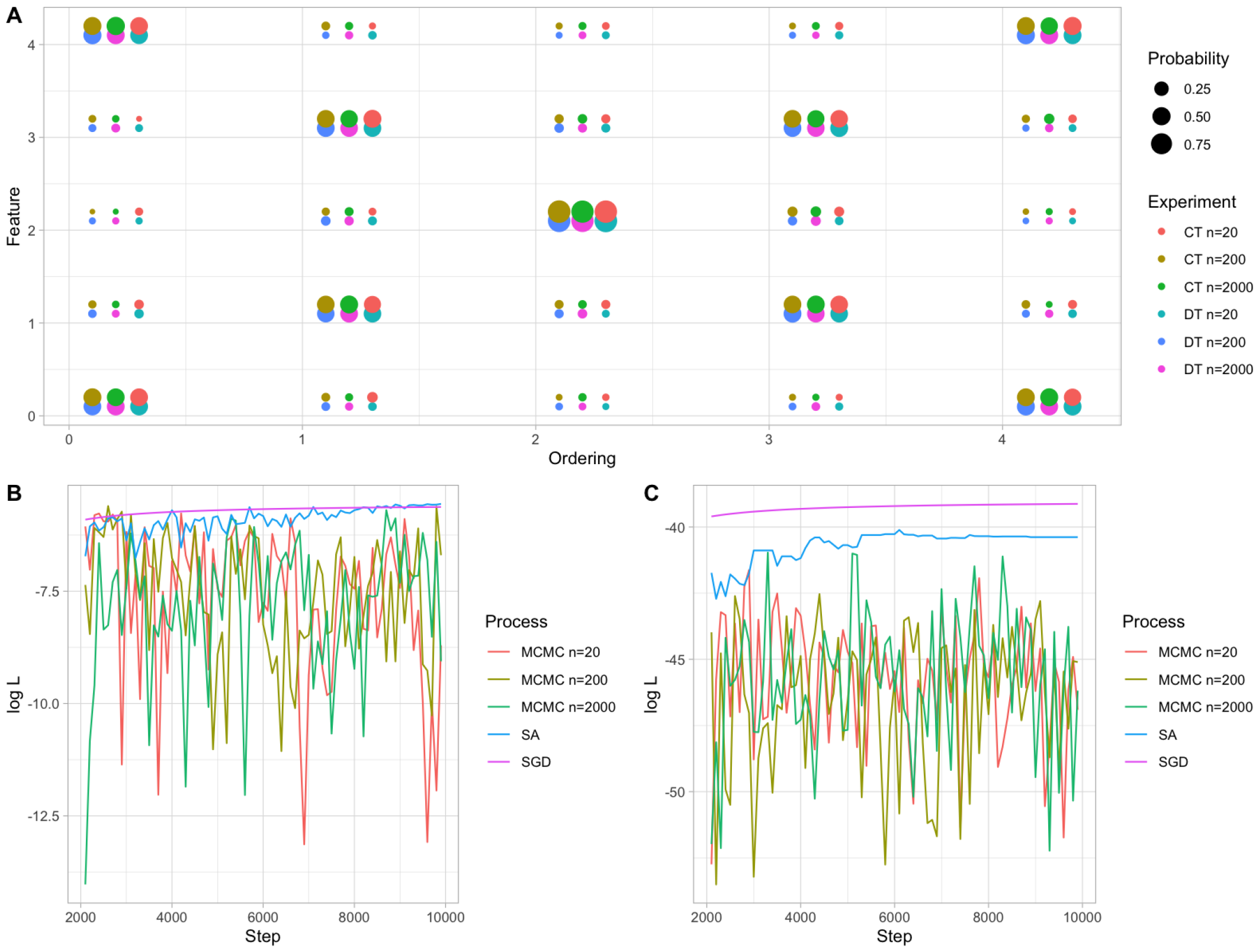
Continuous and discrete time HyperTraPS inference with different numbers of sampling walkers *N*_*h*_, for the same two-pathway test model as in Supp. Fig. S2. Likelihood throughout algorithm progress for MCMC chains with different numbers of sampling walkers, stochastic gradient descent (SGD), and simulated annealing (SA) optimisers.

**Figure S4:**
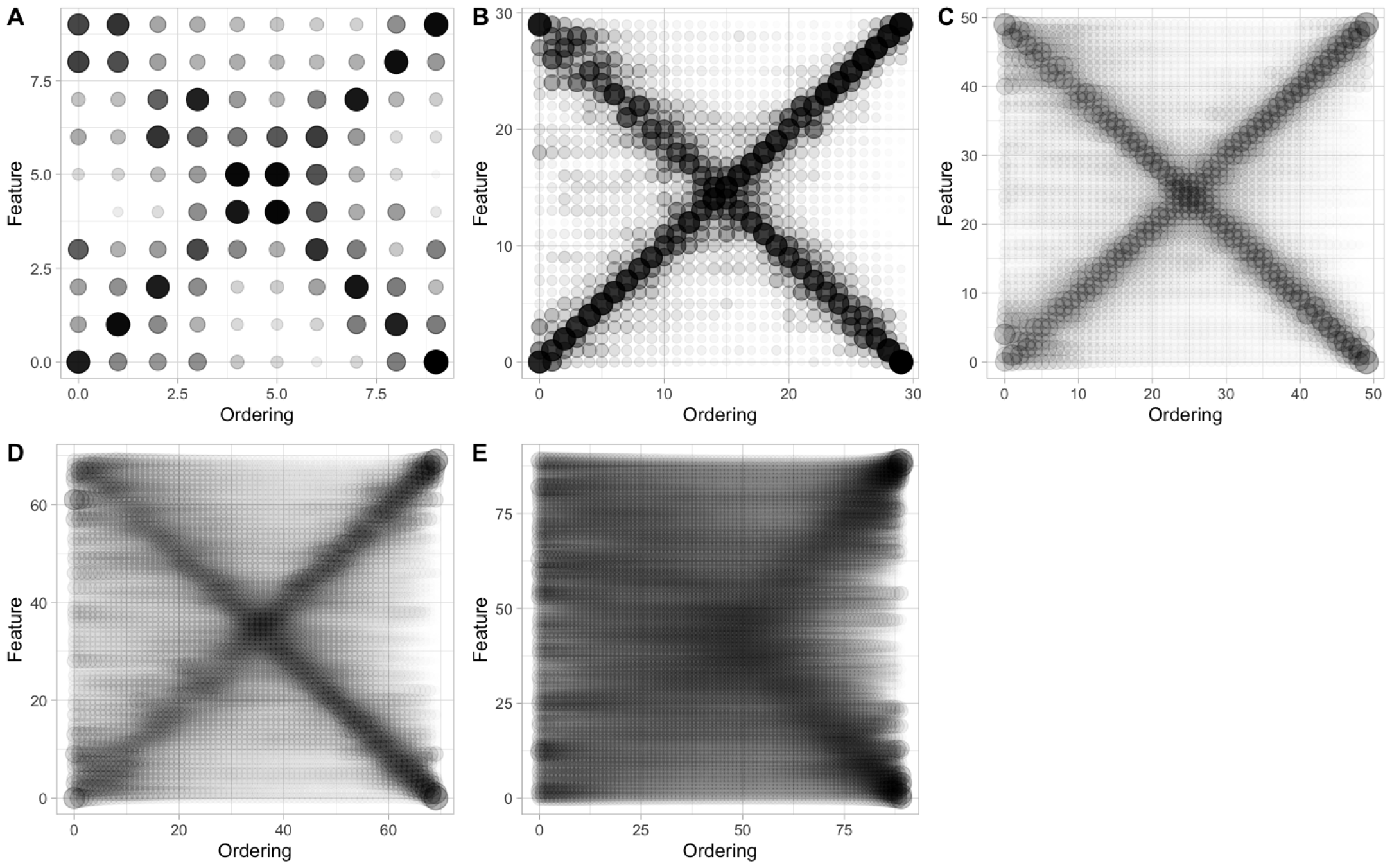
Posteriors on event ordering from discrete time HyperTraPS with increasing numbers of feature *L* = 10, 30, 50, 70, 90. The two-pathway structure generated the data in each case.

**Figure S5:**
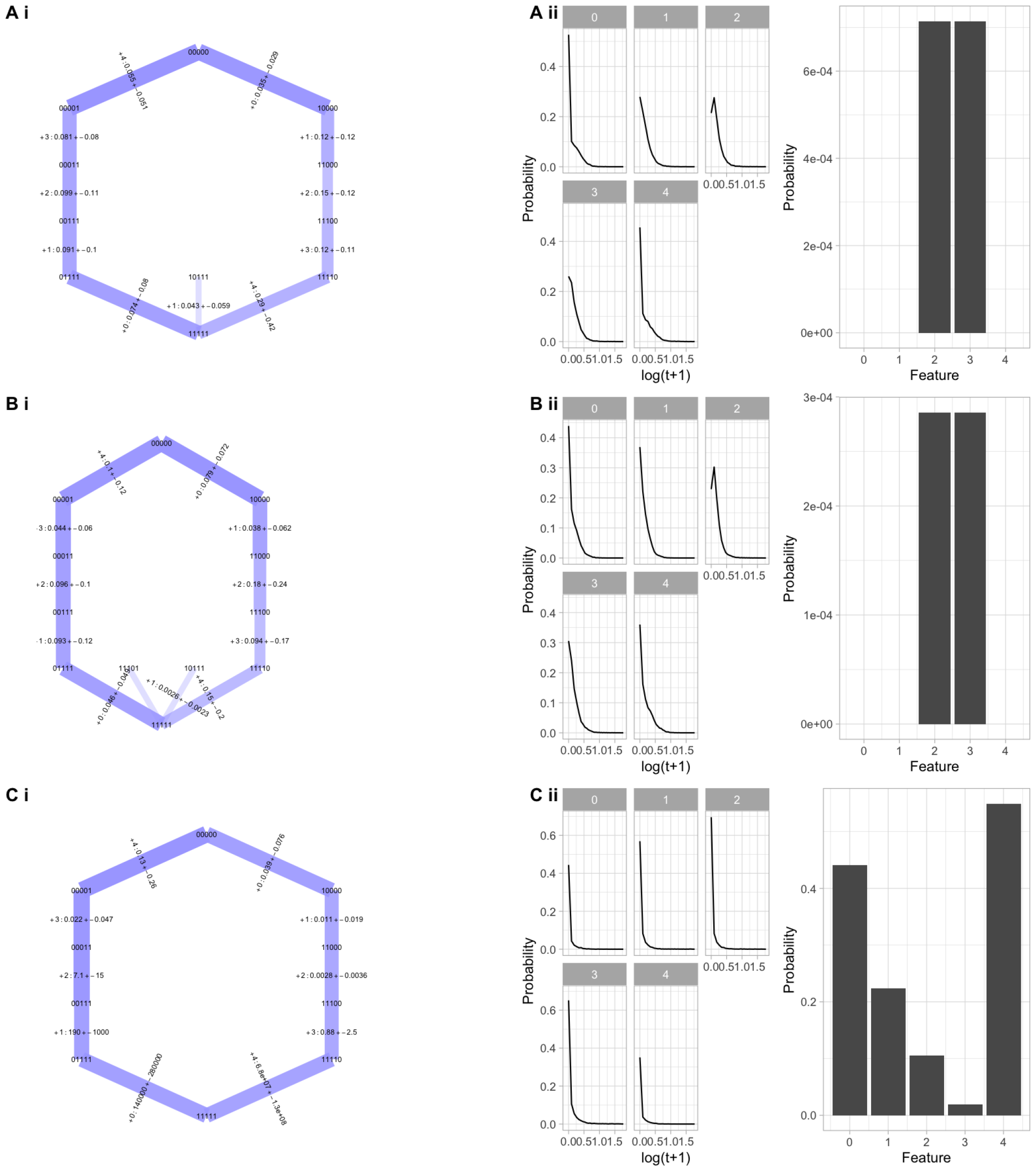
Influence of precision or uncertainty on transition timings. Here, inference is performed using the same illustrative competing-pathway system as throughout the manuscript, but the distribution of timings of each event are specified from different distributions. If the ‘true’ generating process gave timing *τ*_*i*_ for the *i*th observation, these were (A) precisely specified, [*τ*_*i*_, *τ*_*i*_]; (B) uncertain time window, [*τ*_*i*_*/*4, 4*τ*_*i*_], (C) infinite time window, [0, inf]. (C) corresponds to the case where only orderings, not absolute timings, are considered. As previously, (i) give the hypercubic transition networks with mean and s.d. for the timing of each step; (ii) histograms give the distribution of acquisition times associated with each feature, and bar plots give the probability that acquisition does not occur within a threshold time (here, 3 time units).

**Figure S6:**
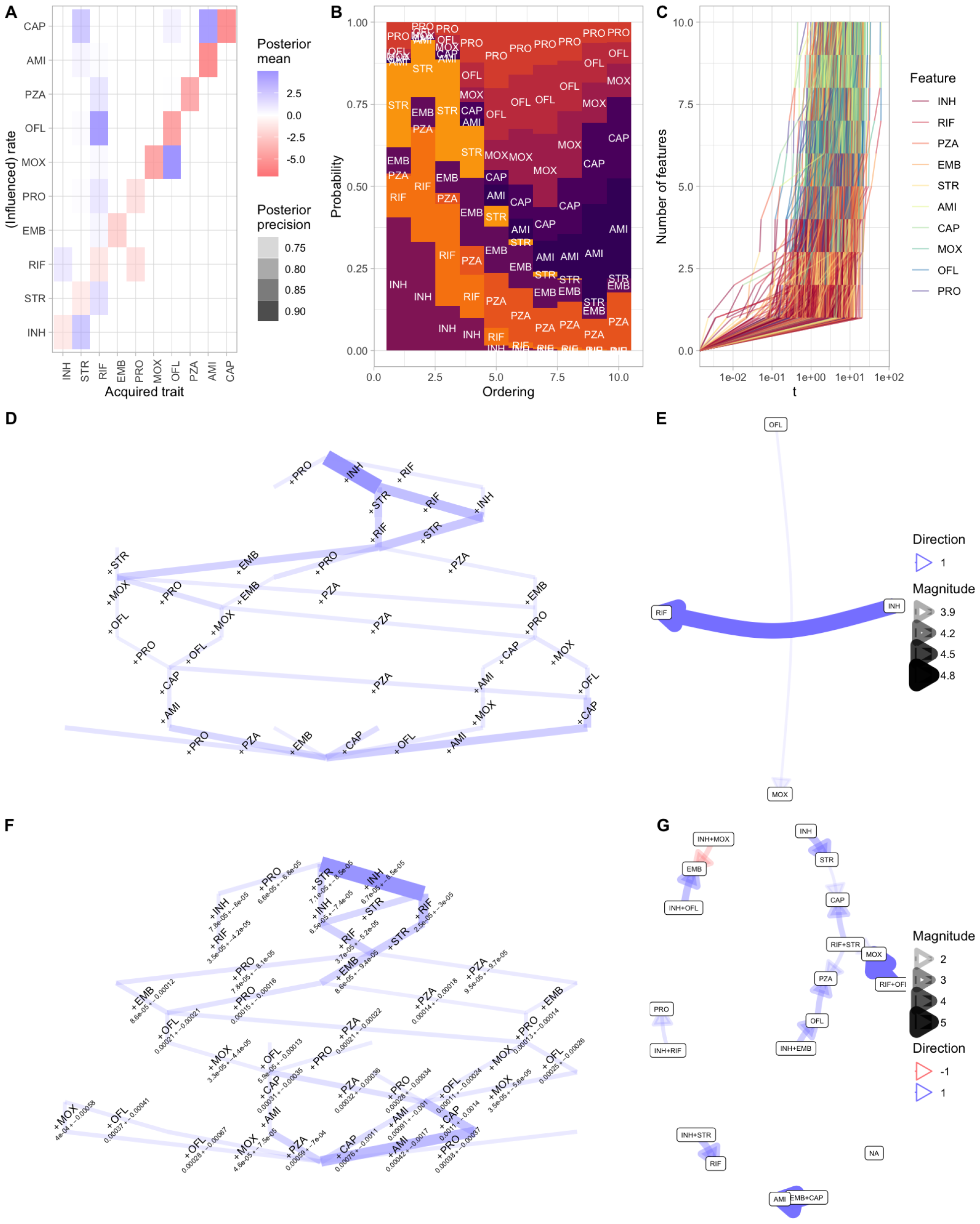
Additional and alternative outputs of inference for the anti-microbial resistance evolution system. (A-C) Additional plots accompanying Fig. 4: (A) Map of inferred influences between drug resistances; (B) Motif plot giving the probability that a given feature is acquired at a particular ordering in the accumulation process; (C) Ensemble of sampled time series of the evolutionary process; here ‘time’ corresponds to the amount of mutational change in the original phylogenetically-embedded data. (D) Inferred transition network for the discrete time case, ignoring timing information, mirroring previous findings (Greenbury et al., 2020) and with less emphasis on early STR (streptomycin) acquisition. (E-F) Output including uncertainty on the branch lengths in the source data phylogeny: (E) more limited network of reliably inferred influence; (F) transition timings with lower modal values. (G) Inferred influences of pairs of features in the *L*^3^ system, thresholded by a posterior coefficient of variation of 0.3.

**Figure S7:**
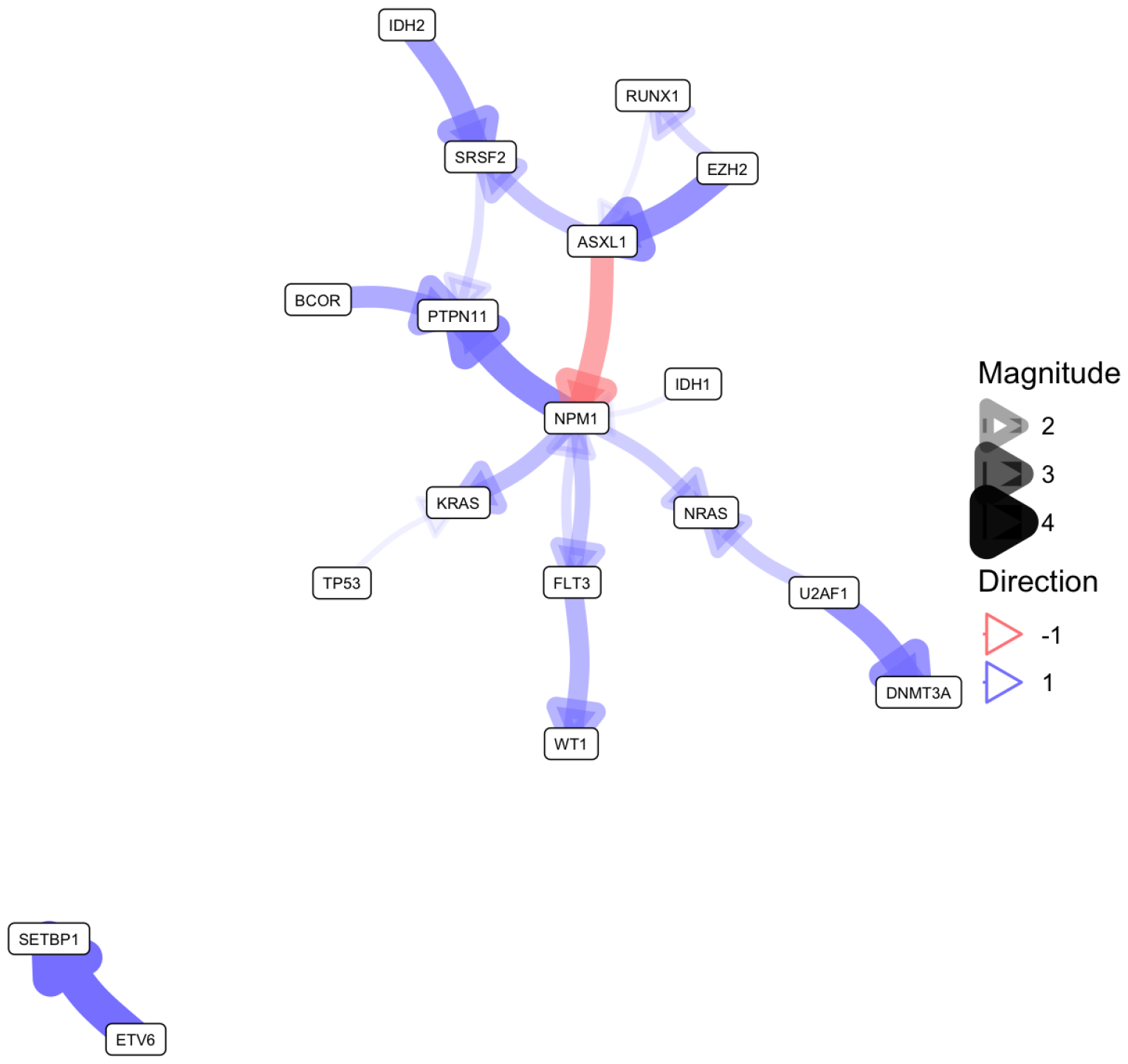
Network visualisation of influences between feature acquisitions in the cancer progression case study.

### Validation, inclusion of prior information, and analysis of posteriors

To test the HyperTraPS-CT algorithm, we first tested the ability of Eqn. 1 and Algorithm 1 to estimate the probabilities underlying accumulation dynamics in continuous time. To this end, we constructed a simple *L* = 3 model system and considered the probability that the system was in state 110 at time *t* = *τ* given that it was in 000 at time *t* = 0, for a set of randomly-chosen parameterisations *λ*. For this simple system, analytic results are readily available and uniform sampling also yields a good estimate of this probability over time. Fig. S2A shows that HyperTraPS-CT (with *N*_*h*_ = 10^3^) readily matches these results, demonstrating the accuracy of the approach in inferring evolutionary timescales.

We next tested the ability of Algorithm 2 to infer parameters underlying evolutionary observations. We constructed a synthetic dataset from a model system with *L* = 3 and a fixed set of transition rates (Fig. S2B). This synthetic system models a case where a slow first transition facilitates a fast second transition, and then a final transition occurs more slowly. We simulated *n* = 10^3^ independent observations at *t*_1_ ∼ U(0, 20), *t*_2_ ∼ *τ*_1_ + U(0, 10) from this system and used these synthetic observations with Algorithm 2 to produce posterior distributions on the underlying parameters. Fig. S2B shows the results of this process, where HyperTraPS-CT readily recovers the dynamic structure of the evolutionary system. We also simulated *n* = 10^3^ independent synthetic observations from an *L* = 3 with a full set of randomly chosen transition rates and *t*_1_ ∼ U(0, 2), *t*_2_ ∼ *t*_1_ + U(0, 5). HyperTraPS-CT recovers the underlying parameters reliably, with few posteriors exhibiting a mode substantially differing from the true value (Figs. S2C, D).

This validation experiment serves to emphasise a principle in the analysis of posteriors produced by HyperTraPS. In some contexts, some processes in the transition network may be very weakly constrained by data – for example, the 010 → 110 and 011 → 111 transitions in Fig. S2B (labelled 2 → 6 and 3 → 7 respectively). In such cases, where systems are highly constrained to follow only a subset of paths, a heuristic can be employed for simplification: set to zero the rates of any processes that are encountered in a proportion lower than *α* of trajectories. For example, *α* = 0.01 would remove all steps that had a less than 1% probability of occurring. As *L* increases, the probability of specific pathways may be expected to decrease entropically, and choices for *α* should be made appropriately. This ‘thresholding’ approach has recently been used to construct a distance metric between inferred hypercubic transition graphs (Garcia Pascual et al., 2024).

We next tested the ability of HyperTraPS-CT to infer distinct evolutionary pathways given observations coupled by a phylogenetic relationship. We considered a larger test case in this instance, by constructing an *L* = 5 systems supporting two competing pathways ‘right-first’ 00000 → 00001 → 00011…, and ‘left-first’ 00000 → 10000 → 11000…. We constructed synthetic datasets of varying size by simulating outputs from this network. Observations were recorded as ancestor-descendant end states, modelling the phylogenetic reconstruction we use throughout this article and in Johnston and Williams (2016).

Figs. S2E-F show the ability of HyperTraPS-CT to reproduce these constrained pathways. The resulting posterior ordering distributions (Fig. S2E) show that HyperTraPS-CT recovers the double pathway structure perfectly for large sets of observations, and very satisfactorily for smaller sets. Fig. S2F shows histograms of the inferred continuous timing of each feature acquisition, well recovering the original timing of 0.1 time units per event.

Fig. S3 shows the behaviour of HyperTraPS and HyperTraPS-CT with different trajectory sample counts *N*_*h*_ and used within different schemes for parameter inference: MCMC for a Bayesian approach, and maximum likelihood via simulated annealing and stochastic gradient descent. Fig. S4 shows the behaviour as the number of features *L* in the dataset is increased; for each *L*, 2*L* observations are produced, so each transition on the hypercube has exactly one associated observation. Even for this limited observation set, the structure of the underlying network is well captured for large feature sets *L*.

## Notes

https://github.com/StochasticBiology/hypertraps-ct

### Competing Interest Statement

The authors have declared no competing interest.

